# Human Interleukin-4-Dependent Facilitation of Human IgG Production in PBL-NOG-hIL-4-Tg mice

**DOI:** 10.1101/2025.10.29.685233

**Authors:** Yoshie Kametani, Shino Ohshima, Ryoji Ito, Yusuke Ohno, Soga Yamada, Yuki Hoshino, Asuka Miyamoto, Mao Suzuki, Nagi Katano, Banri Tsuda, Mariko Miyazawa, Hirofumi Kashiwagi, Daiki Kirigaya, Tomoka Shimizu, Mika Kojima, Yusuke Kikuchi, Shunsuke Nakada, Rentaro Ohki, Atsushi Yasuda, Ayako Hirota, Yukio Nakamura, Jerzy K. Kulski, Tomotaka Mabuchi, Hitoshi Ishimoto, Takashi Shiina

**Affiliations:** Department of Molecular Life Science, Division of Basic Medical Science, Tokai University School of Medicine, Isehara, Japan; Institute of Advanced Biosciences, Tokai University, Kanagawa, Japan; Central Institute for Experimental Medicine and Life Science, Kawasaki, Japan; Department of Palliative Medicine, Tokai University School of Medicine, Isehara, Japan; Department of Obstetrics and Gynecology, Tokai University School of Medicine, Isehara, Japan; Department of Internal Medicine, Division of Nephrology, Endocrinology, and Metabolism, Tokai University School of Medicine, Isehara, Japan; Department of Dermatology, Tokai University School of Medicine, Isehara, Japan; Repertoire Genesis Inc., Osaka, Japan; Microbiology and Immunology, School of Biomedical Science, The University of Western Australia, Crawley, WA, 6009, Australia

**Keywords:** humanized mouse, humoral immunity, T cell receptor, antibody, repertoire, class switch, affinity maturation

## Abstract

Immune-humanized mice provide valuable experimental models for evaluating immune-based therapies, yet the induction of human antigen-specific IgG production remains limited due to species-specific incompatibilities. Our previous work demonstrated that NOG-hIL-4-Tg mice, which express human interleukin-4 (IL-4), support human T and B cell maintenance and enable antigen-specific IgG production following transplantation of human peripheral blood mononuclear cells (PBMCs). In this study, we investigated how IL-4 enhances antibody responses in this model. Flow cytometry and histological analysis revealed that human B cell maintenance was associated with specific plasma hIL-4 concentration ranges at supraphysiological levels. T and B cell receptor repertoire analysis using next-generation sequencing showed that clonal diversity remained largely conserved for one month and decreased to three months post-engraftment. Immunoglobulin repertoire profiling confirmed IgG class switching in an IL-4-concentration-dependent manner. Among the IgG subclasses, IgG3 increased during the first and second months and then decreased thereafter. In contrast, IgG1 tended to increase over time; however, the proportion of IgG subclass varied among individual donors. Following immunization with two distinct peptide antigens, the mice produced enhanced levels of antigen-reactive IgG. However, many B cell clones also exhibited weak responses to unrelated third-party antigens, likely reflecting insufficient affinity maturation. Histological evaluation showed tertiary lymphoid structure (TLS)-like accumulations of B and T cells in the spleen, although fully developed germinal centers were absent. Taken together, these findings demonstrate that NOG-hIL-4-Tg mice maintain human B cells within a regulated IL-4 environment, promote early formation of splenic tertiary lymphoid structures, and support IgG production characterized by clonal expansion, class switching, and somatic hypermutation. These results confirm that human IL-4 expression supports human antibody responses in the PBL-NOG-hIL-4-Tg mouse system.

(278)

## Introduction

Vaccines that induce long-lasting immunological memory play a crucial role in controlling infectious diseases. The rapid development of vaccines against COVID-19 and influenza has been instrumental in mitigating emerging pathogens and enhancing global health security (1, 2). Recent advances in anti-cancer vaccines targeting tumor-specific antigens have also shown promising efficacy (3). Additionally, therapeutic antibodies targeting immune checkpoint molecules, such as PD-1, PD-L1, and CTLA-4, have gained considerable prominence in cancer treatment due to their ability to activate immune responses against cancer cells while reducing the side effects associated with conventional chemotherapeutic agents like cisplatin and methotrexate (4, 5). These immunotherapies enhance the immune system’s capacity to recognize and eliminate cancer cells, offering greater specificity and reduced toxicity. Therapeutic antibodies have also proven effective against infectious diseases, including COVID-19, by selectively targeting pathogens while sparing uninfected cells and tissues (6, 7).

Despite their advances, immune responses to vaccines and immunotherapies vary among individuals due to differences in immune status and genetic background, leading to patient-specific outcomes (8, 9, 10). Comprehensive preclinical data are essential to evaluate the safety and potential side effects of these therapies. However, assessing efficacy remains challenging due to variability and complexity of the immune system across species. To address these challenges, humanized immune mouse models have been developed to better replicate the human immune environment. These models serve as valuable experimental systems for studying humoral immune responses to antigens, antibodies, cells, tissues, and chemotherapeutic agents in preclinical research (11–17).

Transplantation of human hematopoietic stem cells (HSCs), peripheral blood mononuclear cells (PBMCs), or lymphoid tissues into immunodeficient mouse strains is a widely used approach to generate humanized mouse models (18). Among these, PBMC transplantation is often preferred due to the ease of obtaining blood samples and the ability of PBMCs to reflect the donor’s unique immune environment, thereby providing a relatively accurate representation of the human immunity (19, 20). However, a major limitation of PBMC transplantation in immunodeficient mice is the development of graft-versus-host disease (GVHD). Transplanted human T cells can recognize mouse tissues as foreign, triggering a severe xenogeneic immune response that leads to intense inflammation and rapid host death. Under these conditions, the immune system becomes skewed, with a predominance of activated, host-reactive T cells, and a marked reduction of B cells or myeloid cells remaining in lymphoid tissues. To reduce GVHD severity, MHC class I-and class II-deficient mouse strains have been developed to prevent human T cells from recognizing mouse MHC molecules, thereby reducing GVHD severity (21). However, human B cells fail to survive in these MHC-deficient mice, resulting in models that support mainly T cell populations. Although such models are useful for studying T cell function, they are unsuitable for investigating B cell biology.

To overcome this limitation, we developed a humanized (Hu) mouse model, Hu-PBL-NOG-hIL-4-Tg, that produces tumor antigen-specific IgG antibodies following peptide vaccination (22, 23). This model combines human peripheral blood lymphocytes (PBLs) with a NOD/SCID/γc null (NOG) mouse strain genetically engineered to express human interleukin-4 (hIL-4), a cytokine important for modulating Th2 immune responses (24). Our previous studies revealed that B lymphocyte engraftment negatively correlated with glucocorticoid receptor expression in B cells (25), and that immunization with an HER-2 peptide induced antigen-reactive IgG production (14). Additionally, this mouse system distinguished differences in human serum protein composition between healthy donors (HD) and patients with breast cancer (BC) (23) and supported the evaluation of antibody-based anticancer therapies in tumor-bearing humanized mice (26). These findings suggest that the humanized mice system can reflect some of the unique immune characteristics of human donors based on the existence of B cells and antibodies encompassing both cellular and humoral immune responses.

However, the role of IL-4 in facilitating B cell engraftment and antigen-specific IgG production in immunodeficient NOG mice, compared to its role in normal human immune environments remains to be clarified. If the B cell survival rate is high, these mice might maintain a functional immunoglobulin (Ig) repertoire, representing the diversity of B cell receptors (BCRs) capable of responding to antigens. In this hIL-4 environment, antigen-stimulated B cells might undergo clonal expansion and class switch with the help of cognate interaction of T cells. The extent of affinity maturation in engrafted B cells need to be characterized. Affinity maturation is the process by which B cells improve antibody specificity and binding strength through somatic hypermutation (SHM), which introduces point mutations into the variable regions of immunoglobulin (Ig) genes in activated B cells. If affinity maturation is possible, germinal center might be formed within secondary lymphoid organs, where high-affinity B cell clones are selected for survival and differentiation into memory B cells or antibody-secreting plasma cells.

To address these questions, we analyzed the relationship between plasma hIL-4 concentration and the clonal maintenance of transplanted human T and B cells and examined the stability of the Ig repertoire focusing on class switching and SHM, to evaluate the persistence and kinetics of engrafted B cell clones under limited hIL-4 conditions. We also compared IgG responses to HER2 and *Pseudomonas aeruginosa* BamA-derived peptides (27). This analysis provides further insight into the role of human IL-4 in the construction of a human humoral immune system in the mouse environment.

## Materials and Methods

### Ethical approval

This study was conducted in accordance with the guidelines of the Declaration of Helsinki and Japanese federal regulations required for the protection of human participants. The study protocol was approved by the Human Research Committees of Tokai University (12R-002/20R211/21R277) and the Central Institute for Experimental Animals (08-01). Written informed consent was obtained from all the participants. All animal procedures complied with the Guidelines for the Care and Use of Laboratory Animals and were approved by the Animal Care Committees of the Tokai University School of Medicine (174024/185016/191073/202049/213047/ 224039) and the Central Institute for Experimental Animals (20045). This study adhered to the ARRIVE guidelines to ensure rigorous reporting of animal research. Because the experimental design involved constructing a human immune environment in multiple mice using healthy human donor cells, blinding was unnecessary.

### Human peripheral mononuclear cells (PBMCs)

Freshly prepared PBMCs were used in all experiments. Peripheral blood samples (approximately 30 mL) were collected from 75 healthy human donors (32 males and 43 females; age range, 21–64 years) with no history of malignancy, using Vacutainer ACD tubes containing heparin (Becton Dickinson, NJ, USA). PBMC samples were layered over 10 mL of Ficoll-Paque PLUS (Cytiva, London, UK), isolated by density gradient centrifugation (500 ×*g* for 30 min, 20°C), and washed by centrifugation in phosphate-buffered saline (PBS) (300 ×*g* for 5 min, 4°C).

### Mouse strains

NOD/Shi-scid-IL2rγ^null^ (NOG; formal name, NOD.Cg-Prkdc^scid^Il2rg^tm1Sug^/ShiJic) mice or NOG-hIL-4 Tg mice were maintained under specific pathogen-free conditions at the Tokai University School of Medicine. Both mouse strains were healthy, and no obvious defects were observed such as shortened lifespan, smaller number of litters or low body weight. DNA extracted from ear tissue was used for genotyping, and offspring expressing the human IL-4 transgene were identified as described previously (22). Plasma hIL-4 concentrations were measured as described below. NOG mice were obtained from the Central Institute for Experimental Animals, and BALB/c mice were purchased from CLEA Japan Inc. (Tokyo, Japan). All animals were housed under specific pathogen-free conditions at either CLEA or Tokai University.

### Transplantation and treatment of humanized mice

Human PBMCs (5×10^6^ cells) were intravenously injected into 8-week-old NOG or NOG-hIL-4-Tg mice. Antigens emulsified with adjuvant or PBS alone were administered biweekly starting immediately after PBMC transplantation. An overview of the experimental timeline, including transplantation and sample collection time points for various analyses of human adaptive immunity, is shown in **Figure 1A, 2A, 7A.** GVHD was diagnosed based on clinical symptoms and immune profiling. Mice exhibiting hunched posture, ruffled fur, diarrhea, or skin lesions were weighed, and those weighing <20 g were euthanized. Prior to sacrifice, the animals were anesthetized with 4% isoflurane inhalation. Once fully anesthetized, cervical dislocation was performed, and peripheral blood was collected from the heart, followed by organ removal. When possible, the human lymphocyte profiles were analyzed, and GVHD was confirmed if CD3+ T cells exceeded 60%.

**Figure 1.**
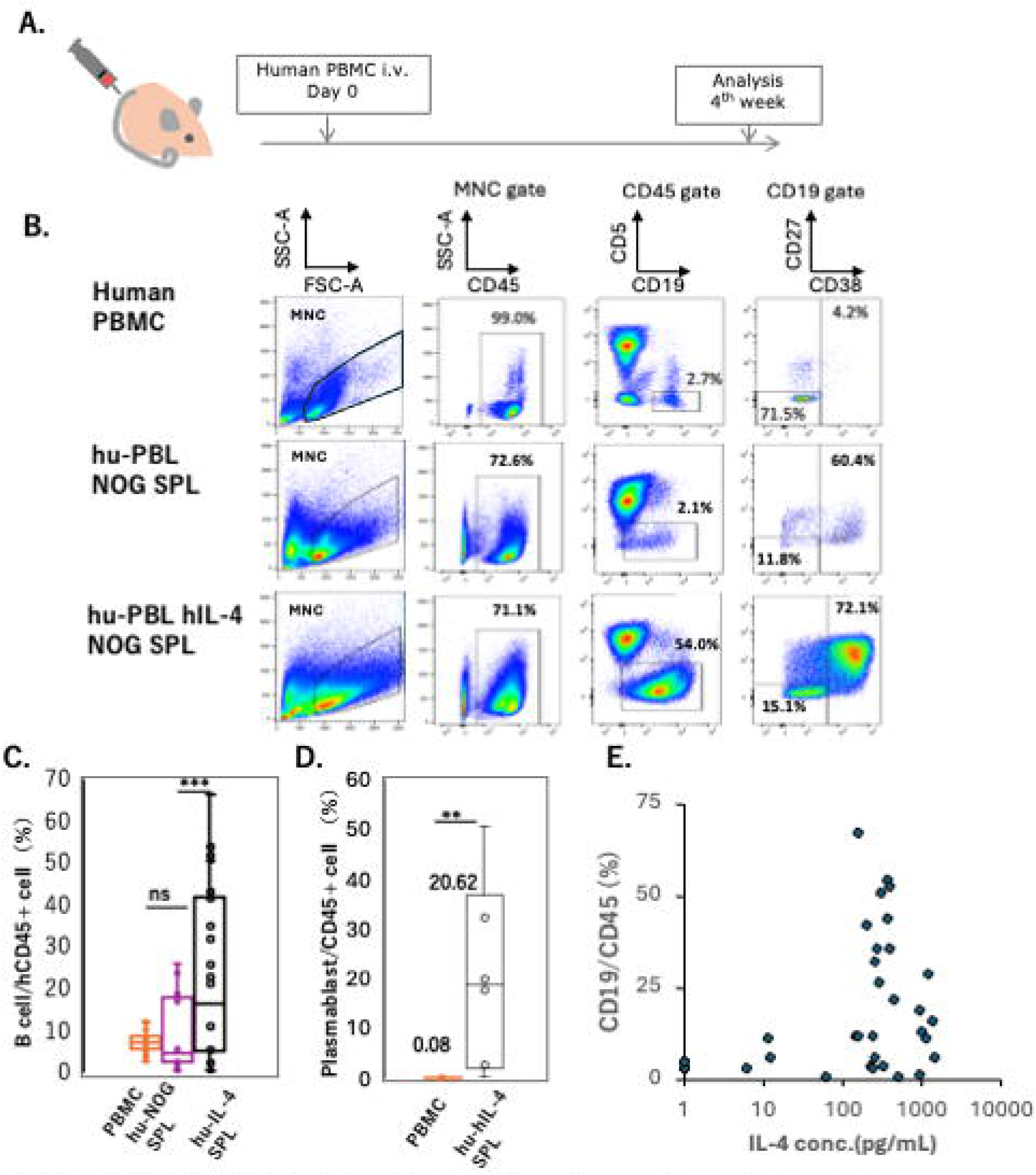
B cell and plasmablast engraftment in the hu-PBL hIL-4 NOG mice. (A) Experimental protocol for the human lymphocyte analysis. (B) Representative flow cytometry (FCM) patterns of original PBMCs (top row), spleen cells from hu-PBL-NOG mice (middle row), and hu-PBL-NOG-hIL-4-Tg mice (bottom row). The percentage of each gated subset is indicated within the panels. MNC gate means mononuclear cell gate. (C, D) Box plots show the percentage of hCD19+ B cells in hCD45+ leukocytes (panel B; PBMCs, n=36; hu-PBL hIL-4 NOG, n=36; hu-PBL NOG, n=16) and the percentage of hCD38+ plasmablasts among hCD45+ cells (panel C, n=8). The horizontal and vertical lines within each box plot represents the mean ± S.D, respectively. Hu-NOG SPL; Hu-PBL-NOG mouse spleen cell, hu-IL-4 SPL; Hu-PBL-NOG-hIL-4-Tg mouse spleen cell. (E) Scatter plot between plasma hIL-4 concentration (X-axis) and the percentage of CD19+ B cells among CD45+ cells (Y-axis). Each dot represents an individual mouse (n=35). Statistical significance was determined using ANOVA and Turkey’s method (B) and Student’s *t*-test (C): p < 0.05 (*) , p < 0.01 (**), and ns (not significant).

### Repertoire analysis of total RNA of engrafted lymphocytes

Total RNA was isolated from 25 samples (4 PBMC samples and 21 mouse spleens; **Supplementary Table 1**) using the RNeasy Plus Universal Mini Kit (Qiagen, Hilden, Germany). RNA concentration and purity were assessed using a NanoDrop Lite Spectrophotometer (Thermo Scientific, Wilmington, DE, USA) and Agilent 2200 TapeStation (Agilent Technologies, Palo Alto, CA, USA). Next-generation sequencing of T-cell receptor (TCR) and B-cell receptor (BCR) repertoires was performed using the unbiased technology developed by Repertoire Genesis Inc. (Osaka, Japan) (28, 29). Double-stranded (ds)-cDNA was synthesized using Escherichia coli DNA polymerase I (Invitrogen), DNA Ligase (Invitrogen), and RNase H (Invitrogen). After end repair with T4 DNA polymerase (Invitrogen), a P10EA/P20EA adaptor was ligated to the 5’ end and digested with *NotI*. Adaptor and primer removal was performed with Agencourt AMPure XP (Beckman Coulter, Brea, CA).

PCR was performed using KAPA HiFi DNA Polymerase (Kapa Biosystems, Woburn, MA) with constant region-specific and P20EA primers (20 cycles: 98 °C for 20 s; 65°C (TCR beta) or 60°C (BCR IgG) for 30 s; 72 °C for 1 min). A second PCR was performed with the same P20EA primers and reaction conditions. The final amplicons were generated using Tag-specific and P22EA-ST1-R primers. Indexing (barcoding) was performed using the IDT for Illumina DNA/RNA UD indexes Set A (Illumina, San Diego, CA).

Amplicons were pooled in equimolar amounts, quantified with a Qubit 2.0 Fluorometer (Thermo Fisher Scientific, Waltham, MA, USA), and sequenced using the Illumina MiSeq platform (2 x 300 bp paired end). Reads were assigned based on identity with reference sequences from the International ImMunoGeneTics Information System (IMGT) database (http://www.imgt.org) using Repertoire Genesis (RG) software. RG includes tools for sequence homology searching (BLATN), gene usage graphics, CDR3 length distribution, and automated aggregation. CDR3 sequences, defined from conserved cysteine (Cys104) of the IMGT to phenylalanine/tryptophan (Phe118/ Trp118), were translated into amino acid sequences. A unique sequence read (USR) was defined by a unique combination of gene segment assignment and CDR3 amino acid sequence. USR copy numbers were ranked per sample. V, D, J, and C gene usage frequencies were calculated.

### B cell clonal RNA sequence analysis

Clonal diversity of unique B cell clones based on total RNA sequences was visualized using the *ggplot2* package in R.

Shannon index was calculated as

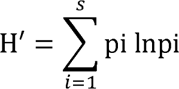

H′ = Shannon index

S = Total number of different species ( Species richness )

Pi = proportion of individuals in the i^th^ species

In = natural logarithm

Class switching from IgM to IgG or IgA was identified by comparing V, D, J, and CDR3 sequences. SHMs were identified by comparing V-region sequences with corresponding germline sequences.

### Flow cytometry

Fluorochrome-conjugated anti-human monoclonal antibodies (mAbs) were used to identify human immune cells in matched PBMC and mouse samples. Cells were incubated with fluorescently labeledmouse anti-human mAbs for 15 min at 4°C, washed with PBS containing 1% (w/v) BSA (Sigma-Aldrich), and analyzed using either a FACS Fortessa or Verse flow cytometer (BD Biosciences, NJ, USA). White blood cells were gated based on the human CD45 expression. Human-specific markers, including CD3, CD5, CD4, CD8, CD11, CD19, CD20, CD27, CD38, CD163, HLA-DR, and PD-1, are listed in Supplementary **Table 2**. Data were analyzed using FlowJo PRIDSCR_008520 software (Becton Dickinson, Franklin Lakes, NJ, USA). Gating strategies are shown in **Supplementary** Figures 1A**, B, and C**.

### Peptide immunization

The CH401 multiple antigen peptide (MAP), which includes the epitope of the anti-HER2 mAb CH401, was used as a cancer antigen (30). CH401MAP was synthesized using Rink amide resin (0.4L0.7 mmol/g) with an ACT357 peptide synthesizer (Advanced Chemtech, Louisville, KY, USA). BAL6MAP, derived from a *Pseudomonas aeruginosa* BAMA peptide sequence (NYYAGGFNSVRGFKDSTLGP), was used as a control antigen (30). CH401MAP or BAL6MAP (50 μg/mouse, 100 μL 1:1 v/v) was emulsified with complete Freund’s Adjuvant (Wako Pure Chemical Industries, Ltd, Osaka, Japan) and administered intraperitoneally into 21 hu-PBL hIL-4 NOG mice and four wildtype BALB/c mice. As a negative control, PBS was emulsified similarly and injected into 36 hu-PBL–hIL-4 NOG mice. A booster immunization using incomplete Freund’s adjuvant was administered two weeks after the first immunization. Mice were sacrificed two weeks after the booster for analysis. Prior to sacrifice, the animals were anesthetized with 4% isoflurane inhalation. Once fully anesthetized, cervical dislocation was performed, and peripheral blood was collected from the heart, followed by organ removal.

### Enzyme-linked immunosorbent assay (ELISA)

hIL-4 protein levels were measured using the Human IL-4 ELISA Set BD OptEIA^TM^ Kit (BD Biosciences), according to the manufacturer’s instructions.

MAP-specific IgG antibody detection was performed as described previously (31). Briefly, micro-wells of microtiter plates (Thermo Fisher Scientific, MA, USA) were coated overnight with MAPs (2 μg/mL) diluted in carbonate buffer (pH 9.5) at 4°C. Wells were washed with PBS-Tween (0.05% v/v) and blocked with 3% BSA-PBS at room temperature (RT) for 1 h. After washing, serial dilutions of mouse plasma were added, and the mixture was incubated at RT for 2 h. Plates were washed, and biotin-conjugated mouse anti-human IgG mAb (Bethyl Laboratories, TX, USA) (1:1000) was added. After incubation for 1 h at RT, the plates were washed with streptavidin-horseradish peroxidase (1:1,000 v/v; Bethyl Laboratories) and incubated for 1 h at RT. Unbound conjugates were removed by washing. Enzyme immunoassay (EIA) substrate kit solution (Bio-Rad Laboratories, Hercules, CA, USA) was added to each well. Anti-human IgG was used to quantify specific antibodies. A known concentration of purified human IgG was prepared, serially diluted, and added as an antigen-mimicking human anti-CH401MAP IgG. A standard curve was plotted based on the absorbance values. As a negative control, the mouse serum was examined using the same protocol. Plasma-specific IgG concentration was calculated from the standard curve. CH401MAP-positive and BAL6MAP-negative clones were counted in the CH401MAP-specific wells.

Hybridoma preparation and characterization of human B cells engrafted in the spleen of the hu-PBL hIL-4 NOG mouse.

Splenocytes from hu-PBL hIL-4 NOG mice or BALB/c mice were fused with the mouse myeloma cell line P3-X63-Ag8-U1, as previously described (32), using electroporation with BEX CFB16-HB evice (BEX Co. LTD, Tokyo, JPN) and electrode LF497P2. Electroporation conditions were set as follows: AC 30 V for 20 s, DC 350 V for 30 µs with a 500 ms DC cycle (repeated three times), followed by AC 30 V for 7 s with fade mode on. Electrofusion was performed in a buffer containing 0.3 M mannitol, 0.1 mM calcium chloride, and 0.1 mM magnesium chloride. Fused cells were cultured in hypoxanthine-aminopterin-thymidine (HAT) selection medium for two weeks. Supernatants were collected and analyzed by LC-MS/MS and ELISA to measure total Ig and CH401MAP-specific IgG levels, as previously described.

### Immunohistochemistry

NOG-hIL-4-Tg mouse spleen, lungs, liver, and tumor tissues were fixed in Mildform® (FUJIFILM Wako Pure Chemical Industries, Ltd.) and embedded in paraffin. Paraffin blocks were sectioned and deparaffinized. Post-fixed tissue sections were stained with hematoxylin and eosin.

For immunohistochemical analysis, tissues from four mice were fixed in formalin, washed, and mounted on glass slides. Endogenous peroxidase activity was blocked for 10 min at RT. Sections were then blocked with goat serum for 30 min, washed, and incubated with primary mAbs as listed in Supplementary Table 2. Subsequently, a peroxidase-labelled anti-mouse IgG antibody (Histofine Simplestain MAX-PO; Nichirei Biosciences INC, Tokyo, Japan) was applied according to the manufacturer’s instructions.

### Statistical analysis

Data are presented as mean ± standard deviation (SD). Statistical significance among groups was evaluated using one-way analysis of variance (ANOVA) and following Turkey’s HSD. As for the comparison of two groups, two-sided Student’s *t*-test was performed. As for the correlation analyses, regression analysis was performed with significance evaluated by F-value. All statistical analyses were performed in Microsoft Excel (RRID:SCR_016137; Microsoft Corp., Redmond, WA, USA).

## Results

### Cellular composition of engrafted human PBMCs

Human B cells, T cells and dendritic cells (DCs) were engrafted with varying efficiencies into the spleens of 18 NOG and 42 NOG-hIL-4-Tg mice for up to 4 weeks after intravenous PBMC transplantation (22) **(Figure 1A)**. At this time point, human CD19+ B and CD3+ T cells were detected in the spleens of both NOG and NOG-hIL-4-Tg mice **(Figure 1B)**. The proportion of B cells among human CD45+ leukocytes in the lymphoid gate was significantly higher in engrafted NOG mice than in the original PBMCs and the overall engrafted cell population **(Figure 1B, C)**. The mean percentage of plasmablasts in eight NOG-hIL-4-Tg mice reached 20.6%, representing a 250-fold increase compared to the 0.1% observed in the original PBMCs **(Figure 1D)**. Additionally, human B cell engraftment was optimal when plasma hIL-4 concentrations were between 100–1,000 pg/mL, indicating a narrow window of hIL-4 levels required for CD19+ B cell maintenance in the spleens of 35 engrafted humanized mice **(Figure 1E)**.

Regarding T cell composition, the CD4/CD8 ratio was higher in NOG-hIL-4-Tg mice than in NOG mice **(Supplementary** Figure 2A). PD-1 and CD25 expression levels were higher in CD4+ T cells than in CD8+ T cells in NOG-hIL-4-Tg mice. In contrast, in NOG mice, PD-1 expression was lower in CD4+ T cells and higher in CD8+ T cells in NOG mice **(Supplementary** Figures 2A and 3).

Among myeloid-derived cells, a significant population of HLA-DR^high^ CD11c^high^ DCs was observed in the monocyte-gated splenocytes of NOG-hIL-4-Tg mice, whereas CD163+ macrophages were absent. While most monocyte-gated splenocytes expressed low levels of HLA-DR and CD11c, the proportion of HLA-DR^high^ CD11c^high^ DCs was markedly lower in four NOG mice (6.2%) than in the 8 NOG-hIL-4-Tg mice (10.3%) **(Supplementary** Figure 2B and C).

CD3LCD56L NK cells were occasionally detected in the spleen; however, their proportion was low, and NK cell engraftment was insufficient in this mouse system (data not shown).

### Kinetics and duration of engrafted human T and B cells

The engraftment dynamics of human T and B cells were monitored over a 3-month period (**Figure 2)**. The experimental protocol is shown in **Figure 2A** and the list of mice that were analyzed are shown in **Supplementary Table 1**. The total number of splenocytes and the proportion of human CD45+ leukocytes peaked during the second month **(Figure 2B and 2C**). T cell numbers and proportions increased over time, eventually comprising the majority of CD45+ cells by the second month **(Figure 2B and 2C**).

**Figure 2.**
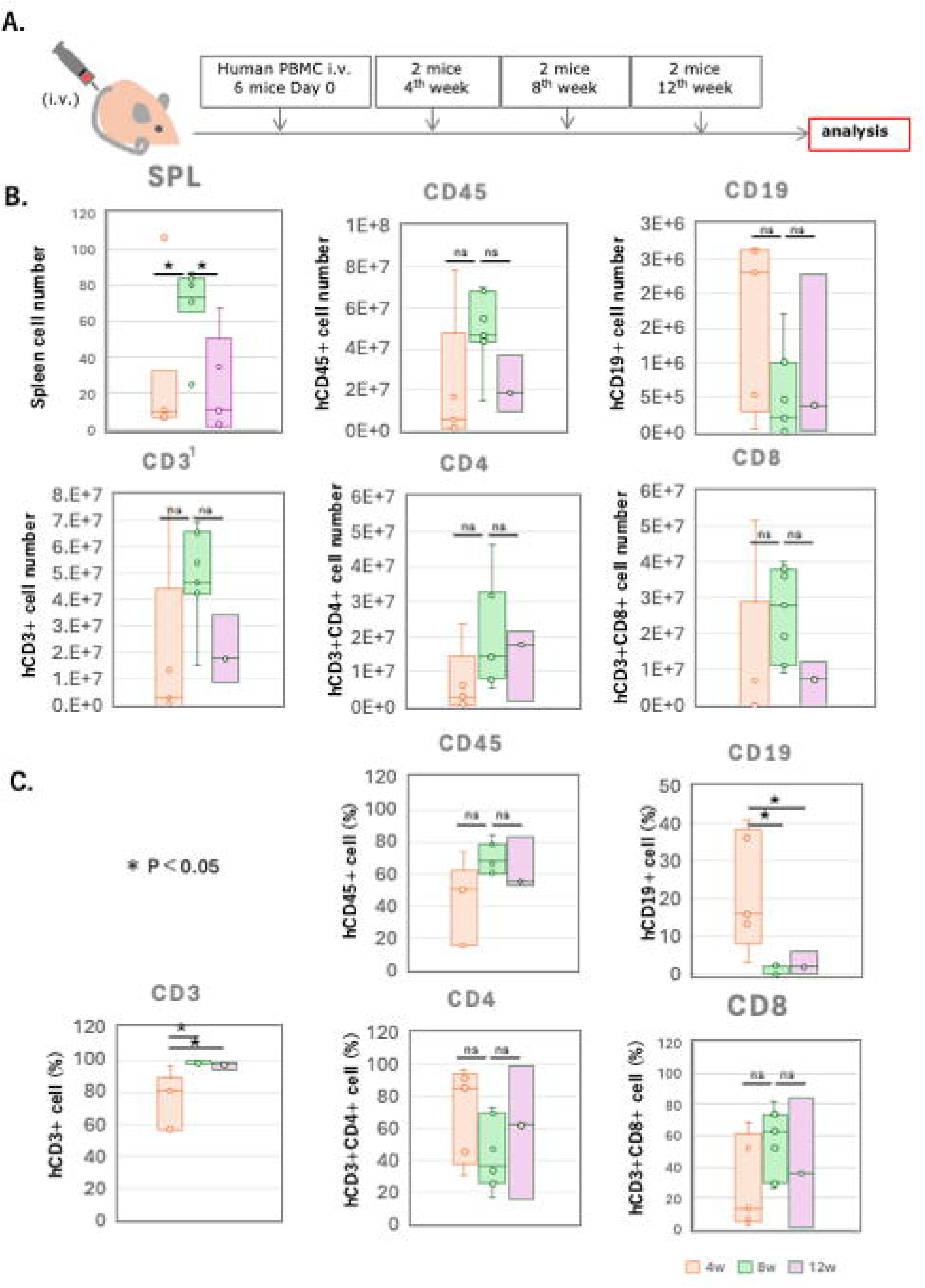
Kinetics of lymphocyte engraftment in NOG-hIL-4-Tg mice. (A) Experimental protocol for the analysis of lymphocyte kinetics. Human PBMCs from each donor (n=4) were divided into 6 aliquots and transplanted into NOG-hIL-4-Tg mice. Two mice were analysed every 4 weeks. Three of the six mice had plasma hIL-4 levels below 100 pmole/L and were excluded from the analysis shown in (B) and (C). (B) Total spleen cell count (upper left panel), hCD45+ human leukocytes (upper middle panel), CD19+/hCD45+ B cells (upper right panel), CD3+/hCD45+ T cells (lower left panel), CD3+CD4+ T helper (Th) cells (lower middle panel), and CD3+CD8+ cytotoxic (Tc) cells (lower right panel). Gating strategy is shown in the Supplemental Figure 1A. (C) Percentage of labelled cell populations over time, measured at 1, 2, and 3 months (Ms) after PBMC transplantation (n=7 for 1M, n=7 for 2M, n=3 for 3M). TIK107 and TIK113 were excluded due to insufficient cell numbers for flow cytometry analysis. Data are presented as mean ± S.D. Statistical significance was determined using ANOVA and thereafter Turkeys Method: p < 0.05 (*), ns (not significant). Data for of individual mice are shown as dots in each box plot. The lines in the box plot indicate the mean score. Detailed sample data are provided in Supplementary Table 1.

CD8+ T cells predominated in the second month but declined in the third month. In contrast, B cells were most abundant in the first month, followed by a decline in the second month, after which their levels remained relatively stable. The expression of CD25+ (an activation marker for conventional and regulatory T cells (Tregs)) and PD-1 (a marker of T cell exhaustion), increased throughout the 3-month period in all mouse groups **(Supplementary** Figure 3). Notably, CD8+ T cells expressed lower levels of these markers compared to CD4+ T cells, suggesting that fewer CD8+ T cells underwent activation-induced exhaustion.

Collectively, these results indicate that both human T and B cell subsets were sustained at significant levels for approximately 2 months, making this timeframe optimal for assessing immune responses, including both humoral and cellular immunity.

### hIL-4 plasma levels and GVHD in humanized mice

Our previous work revealed that most NOG-hIL-4-Tg mice exhibited minimal signs of GVHD after human PBMC transplantation, with symptoms typically emerging 4–8 weeks post-transplantation. We therefore hypothesized that hIL-4 may have a protective role against GVHD. To investigate this hypothesis, we analyzed plasma hIL-4 levels in 24 humanized mice over a 12-week period to determine whether hIL-4 levels influence GVHD onset of severity. **Supplementary Table 1** indicates the GVHD status in these mice, including in 3 representative mice with low hIL-4 levels. Mouse TIK110 (week 4) was diagnosed with GVHD and had a high splenic cell count. Mouse TIK112 (week 8) also developed GVHD, but with a low spleen cell count. Mouse TIK114 succumbed to GVHD before the 12th week. In TIK110, which had low plasma hIL-4 levels, the majority of CD45+ cells were T cells, with a predominance of CD8+ T cells.

Among mice with high hIL-4 plasma levels, 19 of 24 did not develop significant GVHD or showed delayed onset of symptoms. However, two mice (TIK121 and TIK128) with high hIL-4 levels still succumbed to GVHD. These results suggest that while elevated hIL-4 levels may delay GVHD onset or mitigate its severity in many cases, they are not sufficient to completely prevent GVHD in humanized mice.

### Human T and B cell receptor repertoires in PBMC-engrafted mice

To assess the potential for adaptive immune responses in PBL-NOG-hIL-4-Tg mice, we analyzed the TCR and BCR repertoires of the mice shown in **Supplementary Table 1.**

We analyzed the repertoires of original PBMCs and spleen cells at 1, 2, and 3 months after the engraftment of T and B cells in NOG-hIL-4-Tg mice (**Supplementary** Figure 4A**, B**). The TCR and BCR repertoires of the donors were highly diverse, with several clones significantly expanded, suggesting prior exposure of the donors to certain antigens before blood collection. During engraftment in the mice, different clones tended to expand, and no common clone expansion was observed (**Supplementary** Figure 4C). A comparison between IL-4^high^ and IL-4^low^ mice revealed no differences between the two groups. The TCR repertoire remained highly diverse at the end of the first month because the Shannon diversity index maintained approximately two-thirds of its initial value (**Figure 3)**. Moreover, the scores of IgG-producing B cell clone remained stable during the first month, with a significant decrease observed only after 2 months, as reflected in the Shannon diversity index and clone number **(Figure 3A, B)**.

**Figure 3.**
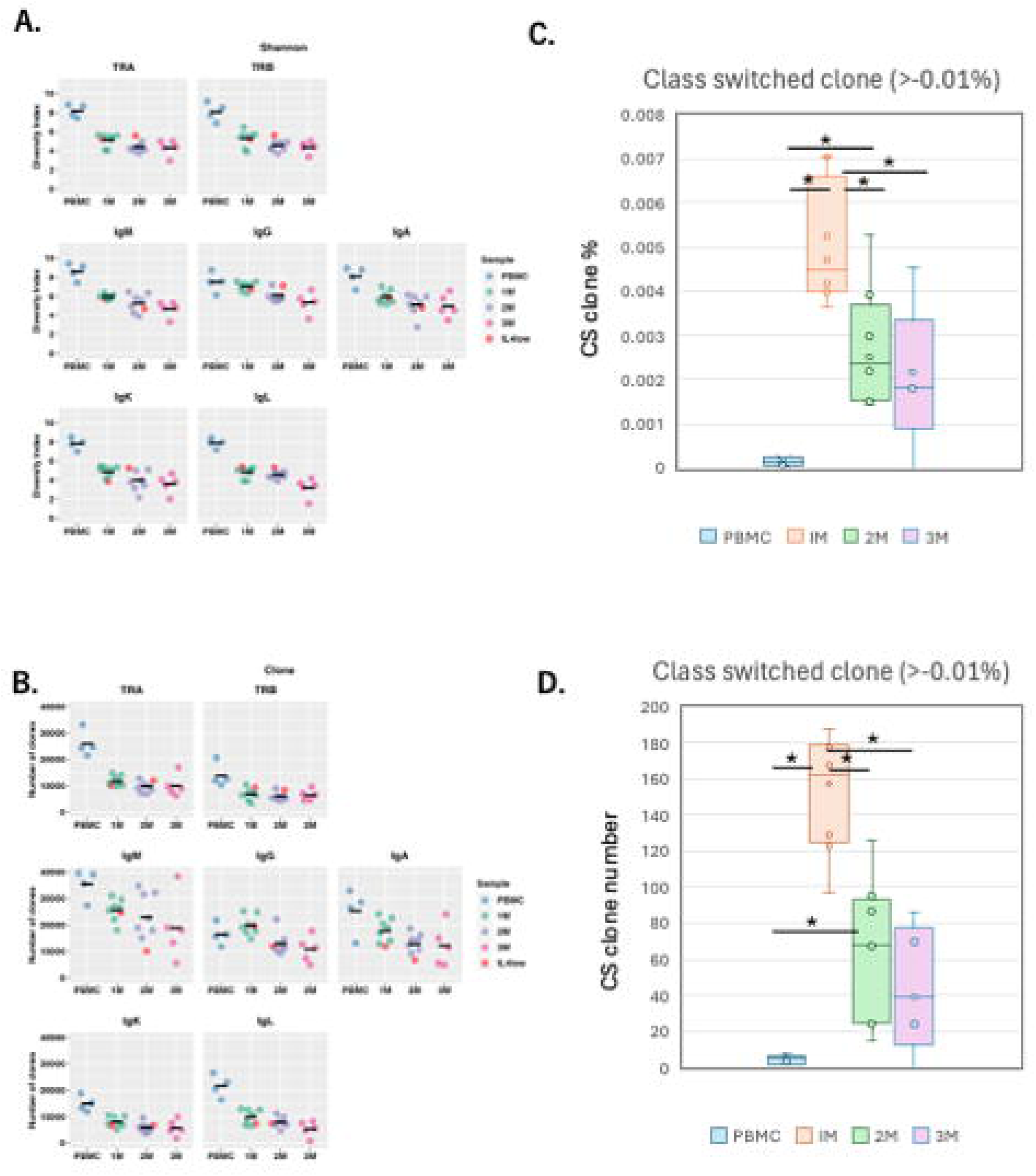
Lymphocyte repertoires in humanized NOG-hIL-4-Tg mice. (A) Shannon diversity Index. Each dot represents the average of Shannon (diversity) index (quantified TCR or BCR diversity) for all analyzed genes examined in an individual mouse at each time point. Orange dots indicate mice with low IL-4 levels. PBMC; n=4, 1M; n=8, 2M; n=8, 3M; n=5. (B) Clone number; Each dot represents the average number of unique reads for all analyzed genes examined in an individual mouse at each time point. PBMC; n=4, 1M; n=8, 2M; n=8, 3M; n=5. The detail of statistical analysis was shown in Supplemental Table 3 and Supplemental Table 4. (C) Box plot of class-switched clone percentage (the proportion of the clone was higher than 0.01%). PBMC; n=4, 1M; n=8, 2M; n=8, 3M; n=5. (D) Box plot of class-switched clone number (the3M). Data for of individual mice are shown as dots in each box plot. The lines in the box plot indicate the mean score. Data are presented as mean ± S.D. Statistical significance was determined ANOVA and thereafter Turkeys method: p < 0.05 (*). Detailed sample data are provided in Supplementary Table 1.

When we examined each mouse data, all mice examined after 1 month retained a substantial number of TCR and BCR clones (**Supplementary Table 3)**. The clone counts gradually decreased in both the TCR and IgG repertoires from the first to third month, to approximately 50% of the original counts in the pre-engrafted PBMCs.

### Ig class switching and somatic hyper mutation

Next, we analyzed whether class switching occurred in the mouse B cells of the individuals shown in **Supplementary Table 1**. Compared to the limited PBMC Ig repertoires before transplantation, both the proportion and the number of class-switched clones increased significantly one month after transplantation and declined thereafter (**Figure 3C, D**). Representative examples of class switching are presented in **Supplementary Table 4**. While most class-switched clones were not detected in the original PBMCs of RG1 (TIK101), as indicated by a percentage of zero, the IgM clones observed in TIK102 (after 1 month) shared identical VDJ rearrangements and CDR3 sequences with IgG or IgA clones.

We then examined the kinetics of clone frequency and the number of IgG subclasses over a three-month period. The frequency of IgG3 clones significantly increased one month after transplantation, followed by a gradual decline after the second month (**Figure 4A**) in 3 out of 4 donors. In contrast, IgG1 clones increased up to the third month in two donors, while the other two showed a decline over the same period. IgG2 and IgG4 did not exhibit any consistent trends. Meanwhile, analysis of the class-switched clone numbers indicated that IgG1 and IgG3 clones were maintained in the mice (**Figure 4B**).

**Figure 4.**
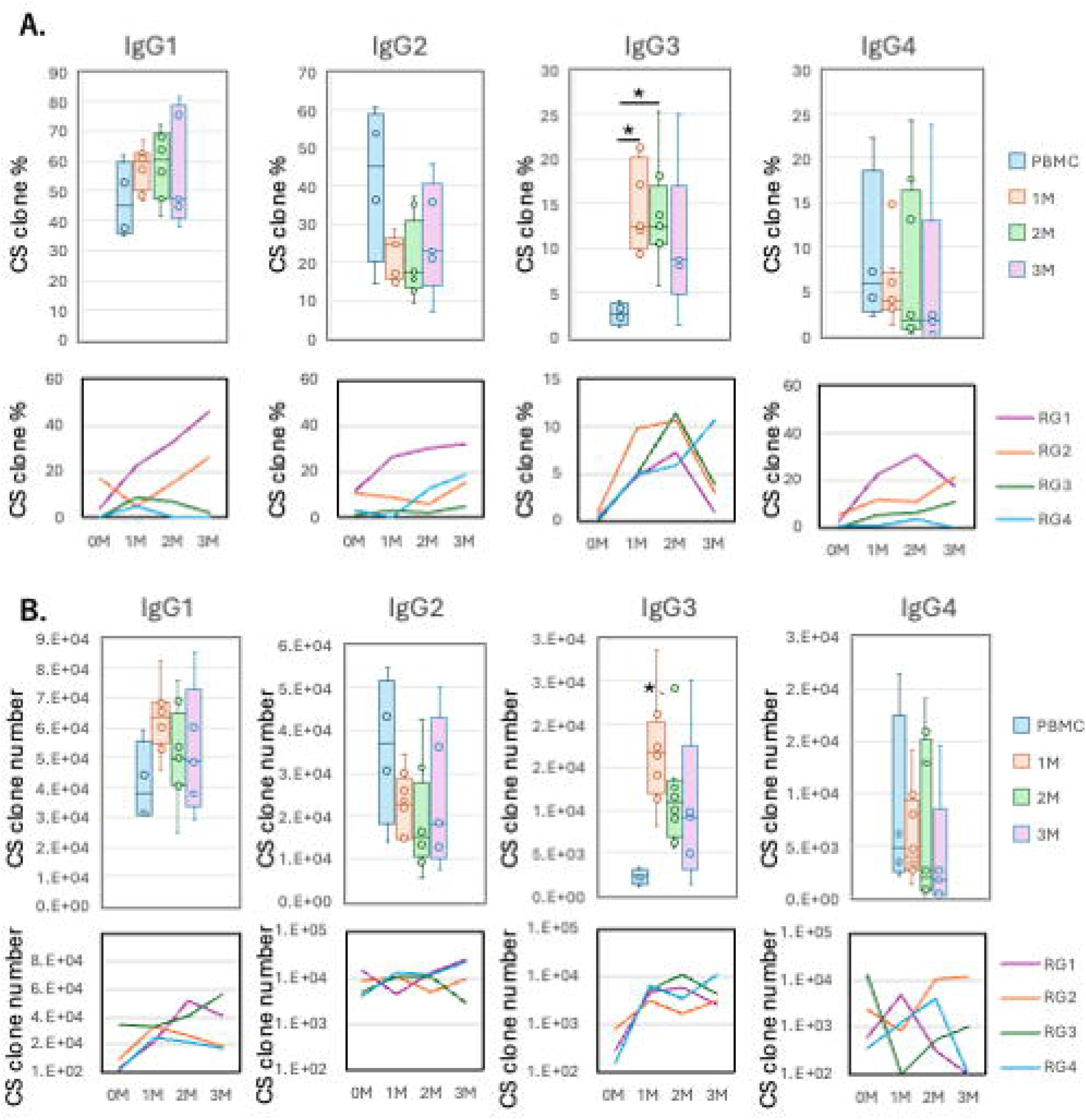
Kinetics of IgG subclass in humanized NOG-hIL-4-Tg mice. (A) Box plots (Upper panels) and lines of each donor’s mean scores (Lower panels) show the percentage of class-switched clones of each IgG subclass (the proportion of the clone was higher than 0.01%). (n=4 for PBMC, n=8 for 1M, n=8 for 2M, n=5 for 3M). RG1-RG4 are the ID of donors (Table 1). (B) Box plot (Upper panels) and lines of each donor’s mean scores (Lower panels) show the number of class-switched clones of each IgG subclass (the proportion of the clone was higher than 0.01%). (n=4 for PBMC, n=8 for 1M, n=8 for 2M, n=5 for 3M). The horizontal and vertical lines within each box plot represents the mean ± S.D, respectively. Individual data were also shown in the box. Statistical significance was determined by ANOVA and thereafter Turkeys method: p < 0.05 (*). Data for of individual mice are shown as dots in each box plot. The lines in the box plot indicate the mean score.Detailed sample data are provided in Supplementary Table 1.

The frequency and distribution of SHM observed in these repertoires are shown in **Figures 5A and 5B**. Compared to the reference genome, SHM was detected at higher frequencies in both PBMCs and engrafted mice **(Figure 5A)**. **Figure 5B** highlights an example from mouse TIK118 (clone #711), where seven amino acid substitutions were identified within the variable region of the Ig heavy chain.

**Figure 5.**
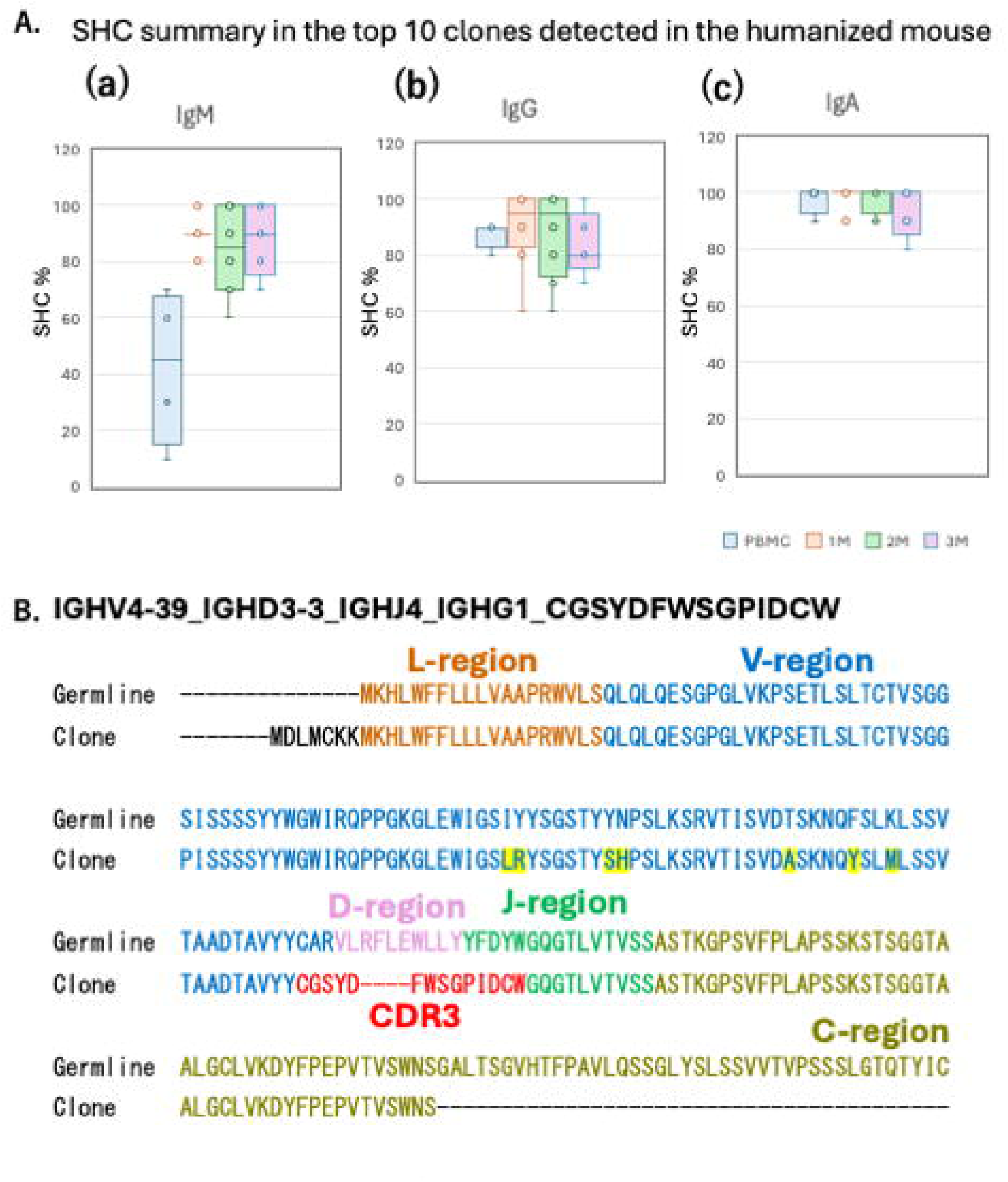
Kinetics of IgG subclass in humanized NOG-hIL-4-Tg mice. (A) Box plot of SHMs detected in the top 10 clones from each of the 21 humanized mice and 4 PBMC analysed for IgM, IgG, and IgA. Data are presented as mean ± S.D. Statistical significance was determined ANOVA and thereafter Turkeys method: p < 0.05 (*). Data for of individual mice are shown as dots in each box plot. The lines in the box plot indicate the mean score. (B) Representative SHM in the IgG heavy chain (mouse TIK118). Germline (upper) and clone (lower) amino acid sequences are aligned, with each region highlighted in a different colour. SHMs are marked in yellow.

To assess whether IL-4 concentration influenced Ig variability, we analyzed correlation between plasma hIL-4 levels and several metrics: Shannon diversity index, specific clone number (unique sequence reads), class switching scores, total class-switching events and subclass-switching events. Plasma hIL-4 concentrations ranged from <500 pg/mL to >100 pg/mL. Data points outside this range showed excessive divergence and were excluded from analysis. As shown in **Figure 6**, nearly all the datasets exhibited a positive linear correlation. Notably, the clone number and IgG class switching were significantly associated with plasma hIL-4 concentration (R²=0.3353, F=0.0118) **(Figure 6C)**. Additionally, the total number of class-switching events strongly correlated with hIL-4 levels (R²=0.5629, F=0.0003349) **(Figure 6A)**. We also observed correlation between IgG1 clone number and IL-4 concentration, and IgG3 tended to be positively correlated with IL-4 concentration during the first 2 months, although the F-value was 0.0534 (**Supplementary** Figure 5). Collectively, these findings indicate that the mice developed substantial variability in their T and B cell repertoires, and that this variability, including the extent of class switching to IgG and SHM, was significantly influenced by plasma hIL-4 levels.

**Figure 6.**
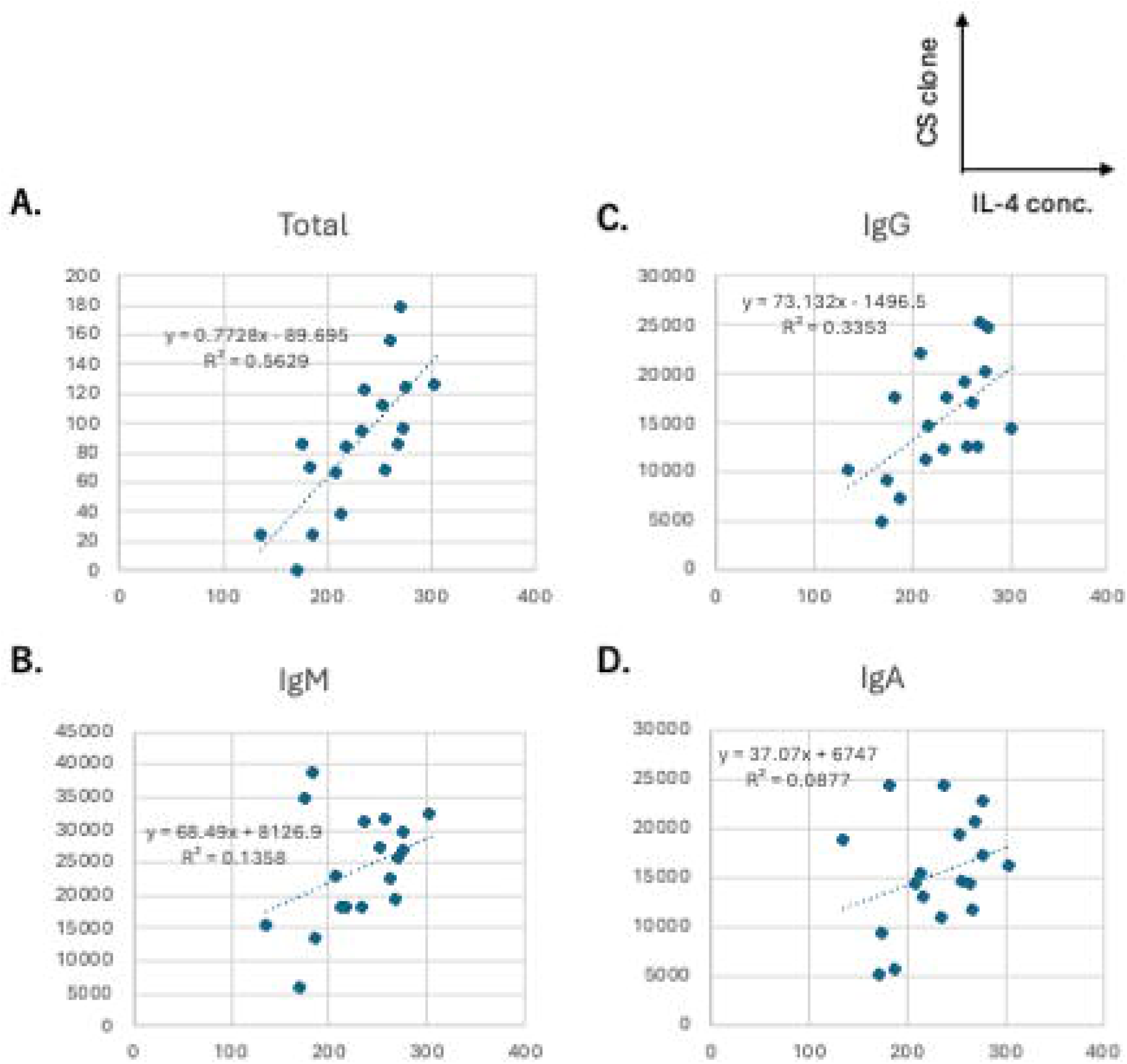
Correlation between IL-4 concentration and the number of class-switched clones. (A) Total clones (>0.01% of total), (B) IgM, (C) IgG, (D) IgA. (n = 18). Samples with low IL-4 levels (<100 pg/mL; TIK110 and TIK112) and high IL-4 levels (>500 pg/mL; TIK105) were excluded. The regression lines, formulas, and R² values are displayed in each panel. F-values for each marker: Total clones = 0.0003349, IgM = 0.1323, IgG = 0.0118, IgA = 0.2329.

### Antigen specificity of human B cell clones in the mice following CH401MAP/ BAL6MAP immunization

While SHM was frequently observed in the PBL-hIL-4-Tg NOG mice, it does not guarantee the occurrence of affinity maturation, by which high-affinity clones are selected and expand. To determine whether affinity maturation could occur in this environment, we assessed the antigen specificity and cross reactivity of B cell clones produced after peptide immunization.

PBL-hIL-4-Tg NOG mice were immunized with either CH401MAP, a multiple aligned peptide (MAP) derived from a 20-mer HER2 epitope, or BAL6MAP, a third-party MAP antigen corresponding to the *Pseudomonas aeruginosa* BAM antigen **(Figure 7A)** (29). As controls, additional mice received adjuvant emulsified in PBS. After booster immunization, mice were sacrificed, sera were collected, and splenocytes were fused with the mouse myeloma cell line P3X to generate hybridomas, following established protocols (32). Antibody titration and hybridoma screening were performed using both CH401MAP and BAL6MAP peptides.

**Figure 7.**
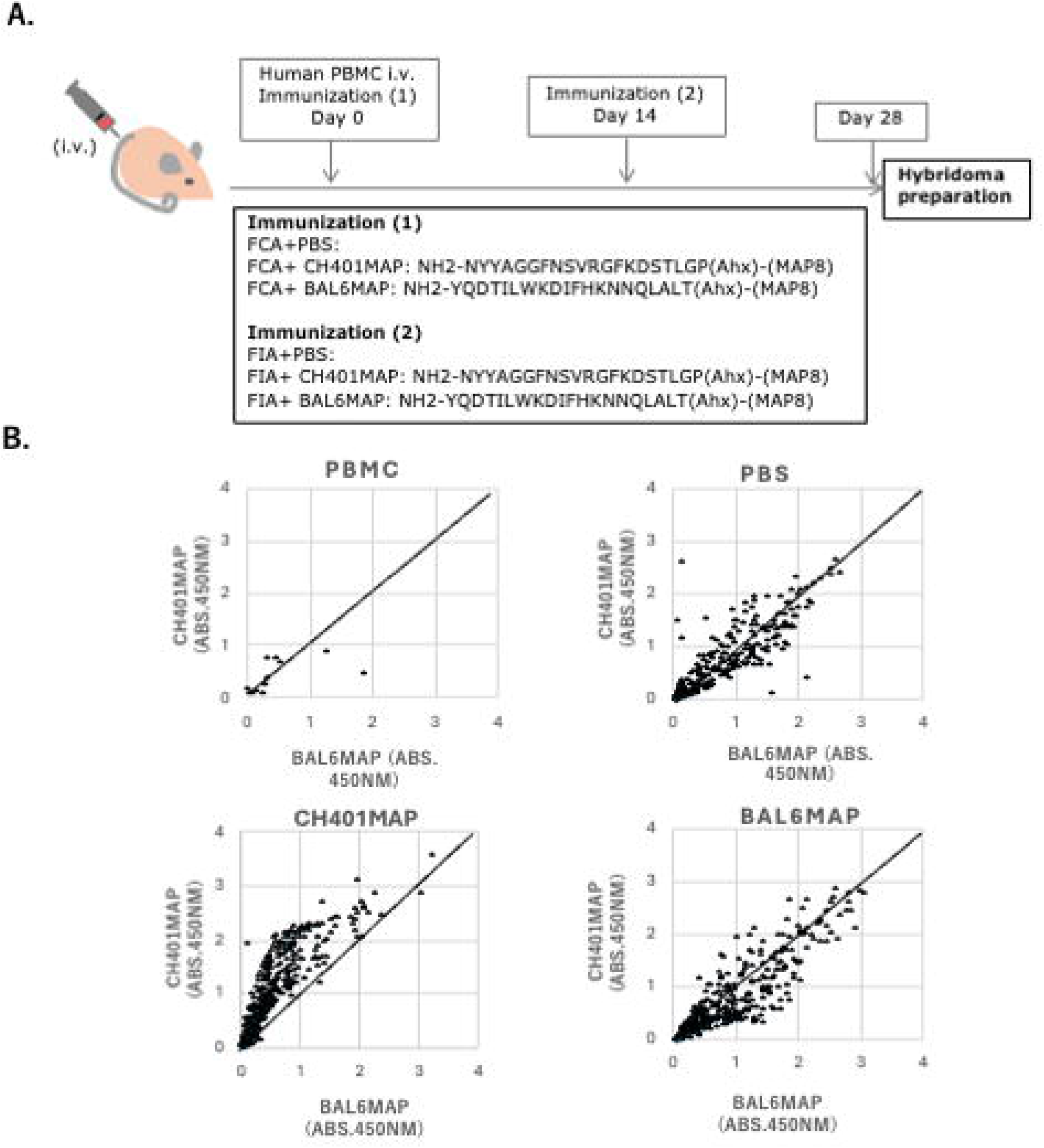
Antigen-specific hybridoma produced from hu-PBL hIL-4 NOG mouse spleen cells. **(A)** Immunization protocol for hu-PBL-NOG-hIL-4-Tg mice. The immunogens used for the primary immunization and booster immunizations are indicated along the timeline. FCA: Freund’s complete adjuvant, FIA: Freund’s incomplete adjuvant. **(B)** Scatter plots showing representative reactivity and specificity of antibodies produced by B cell hybridomas derived from human PBMCs (n=7, upper left panel), PBS-treated control mice (n=4, upper right panel), CH401MAP-immunized mice (n=9, lower left panel) and BAL6MAP-immunized mice (n=4, lower right panel). The vertical axis represents cross-reactivity to CH401MAP, and the horizontal axis represents cross-reactivity to BAL-6MAP. Absorbance at 450 nm is shown. Each dot represents a single hybridoma clone.

As shown in **Supplementary Table 5**, many mice exhibited serum antibody reactivity to CH401MAP. However, most of these sera also reacted to BAL6MAP, indicating a high frequency of cross-reactive or non-specific antibody responses. Since highly specific antibodies might not be detected in total serum titers, we further analyzed the specificity of individual B cell clones by generating hybridomas from splenic B cells.

Hybridomas secreting human antigen-specific IgG were successfully derived from spleen cells of NOG-hIL-4-Tg, whereas only a few hybridoma clones were recovered from the original human PBMCs **(Figure 7B).** In adjuvant-only (PBS)-immunized mice, most hybridoma clones cross-reacted with both CH401MAP and BAL6MAP, suggesting the presence of non-specific or polyreactive clones. In contrast, CH401MAP-immunized mice generated many clones that specifically reacted with CH401MAP, although many of these also exhibited weak cross-reactivity to BAL6MAP **(Figure 7B)**. Several clones demonstrated high specificity for CH401MAP with minimal cross reactivity to the third-party antigen. A similar pattern was observed in BAL6MAP-immunized mice; most clones reacted strongly to BAL6MAP, but many also cross-reacted with CH401MAP.

To compare the humanized mouse response to a wildtype mouse immune system, BALB/c mice were also immunized with both MAP antigens. Specificity analysis revealed that CH401MAP-immunized BALB/c mice produced weakly reactive and poorly specific antibodies, whereas BAL6MAP-immunized mice generated strongly reactive antibodies with higher antigen specificity and fewer cross-reacting clones **(Supplementary** Figure 6).

These findings suggest that while humanized NOG-hIL-4-Tg mice can produce antigen-specific human IgG-secreting hybridomas, the frequency of crossreactive clones to the third-party antigens remains high. In contrast, BALB/c mice produce more specific hybridomas, particularly in response to BAL6MAP, highlighting a limitation in the affinity maturation environment of the humanized mouse model.

### Spatial organization of human lymphocytes in the mice spleens

Given the insufficient affinity maturation observed in this humanized mouse system, it is possible that the germinal center reaction was not fully induced. To explore this, we assessed the spatial distribution of human lymphocytes in the spleens of NOG and NOG-hIL-4-Tg mice using immunohistochemistry.

In NOG mice, no well-organized tertiary lymphoid structures (TLSs) (33) or germinal centers were observed **(Figure 8A)**. In contrast, the spleens of NOG-hIL-4-Tg mice showed multiple B cell clusters surrounded by T cells (**Figure 8B**). These structures resembled early TLS-like formations typically observed in mucosal and tumor tissues, suggesting that B cell accumulation in the spleens of NOG-hIL-4 mice contributes to the development of lymphoid-like microenvironments. However, fully mature germinal centers were not observed.

**Figure 8.**
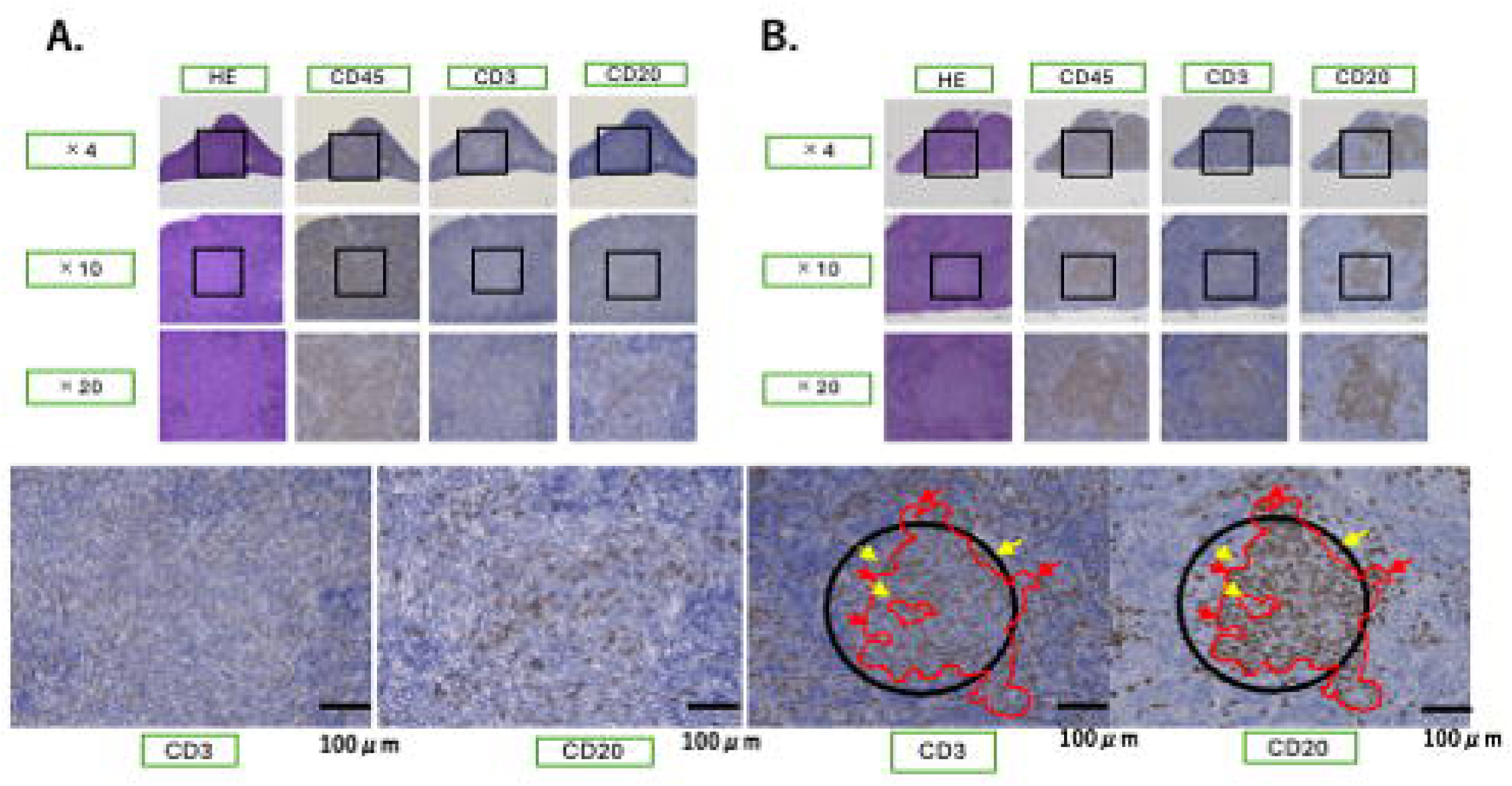
TLS-like structures in the hu-PBL hIL-4 NOG mouse spleen. Representative immunohistochemistry-stained spleen sections from NOG (n=5) (A) and NOG-hIL-4-Tg (n=8) (B) mice. The antibodies used for staining (anti-CD45, anti-CD3, and anti-CD20) are indicated at the top of each panel. Magnification levels are shown on the left. Squares within the images mark the magnified regions (x20) displayed in the lower panels. The four bottom panels highlight the tertiary lymphoid structure (TLS)-like formations in the spleens of NOG-hIL-4-Tg mice. B cells (CD20+) accumulate in clusters, while T cells (CD3+) surround these regions as indicated by black circles. Yellow arrows mark T cell-dominant areas, and red lines indicate B cell-dominant regions. No TLS-like structures were observed in NOG mice.

## Discussion

In previous studies, we demonstrated that humanized immunodeficient NOG-hIL-4-Tg mice successfully support the engraftment of human B and T cells following PBMC transplantation (34), generated tumor antigen-specific IgG antibodies after peptide vaccination (14) and characterized HER2-specific humoral responses in breast cancer-derived lymphocytes (22, 31). However, these studies did not fully address key questions regarding how hIL-4 influences antigen-specific IgG production. To address this issue here, we conducted a comprehensive analysis that included the kinetics of PBMC engraftment, the diversity and stability of human TCR and BCR repertoires, Ig class switching, and SHM. We further assessed the potential for affinity maturation by examining antibody reactivity to unrelated third-party antigens.

Time-course studies revealed that engrafted human B- and T-cell clones expanded and persisted in NOG-hIL-4-Tg mice for up to three months, with B cells peaking during the first month and T cells during the second, followed by a gradual decline in both populations. After transplantation of human PBMCs, proliferating T cells and B cells mostly accumulated in the spleen or bone marrow. In this model, human T cells likely undergo homeostatic proliferation, potentially driven by low-level autoreactive TCR or cytokine signaling, such as IL-7 or CD28-mediated co-stimulation (35, 36). Since hIL-7 was not administered, weak TCR and/or CD28 signaling may have contributed to this proliferation. If CD28 plays a role, its ligands, CD80 and CD86, provided by B cells or DCs in the spleen, may have facilitated the expansion of transplanted PBMCs (5×10^6^ to 2∼10×10^7^ cells). This hypothesis is partially supported by our observation that a high proportion of T cells exhibited a memory phenotype, suggesting a relative absence of quiescent naive T cells.

Although the B cell number increased significantly in the first month, most engrafted human B cells differentiated into plasmablasts, likely due to the inadequacy of the mouse stromal environment needed to support their long-term survival. Moreover, B cell clones underwent class-switching to IgG and/or IgA and acquired SHMs. Notably, class switching to IgG showed a significant positive correlation with plasma IL-4 concentration between 100 pg/mL and 500 pg/mL, suggesting that hIL-4 enhances IgG production in this humanized mouse model. Interestingly, the expansion of IgG3-producing B cell clones preceded that of IgG1-producing B cell clones among the IgG-expressing population. Since the constant region of IgG3 is located proximally to the IgD gene, the earliest class switching may have occurred at the IgG3 region, followed by subsequent class switching to other IgG subclass regions (37,38). The IgG3 region has been reported to contain IL-4-responsive elements (39,40), suggesting that human IL-4 initially induces class switching of the Ig gene to IgG3, with further switching to IgG1 occurring frequently in NOG-hIL-4-Tg mice. Although the distribution of IgG subclass did not correlate with IL-4 concentration, the IL-4 level in these mice appears sufficient to induce such class switching. Mice immunized with CH401MAP or BAL6MAP generated B-cell clones that were cross-reactive with immunizing peptides, although full antigen specificity was not achieved. These findings indicate that a certain degree of clonal selection and expansion occurred within the B-cell repertoire of the humanized mice.

Affinity maturation of B cell clones appeared to be incomplete, likely due to the absence of FDCs and other components essential for germinal center reactions (41). SHM of human Ig genes was observed in the mouse immune system, and tertiary lymphoid structures (TLSs), which represent early-stage structures for affinity maturation (33), were identified in the spleen of humanized mice, where human T cells surrounded clusters of B cells. However, fully developed germinal center-like formations were not observed. This suggests that B cells were unable to continuously receive the antigen stimulation and cytokine signals necessary for the survival and selection of highly specific clones (42). The absence of FDCs may be attributed to the lack of precursor cells within the transplanted PBMCs, potentially explaining why only a limited number of highly specific IgG-secreting clones were detected in the spleens of immunized mice. If PBMCs were sourced from individuals who had recovered from infectious diseases, they might contain pre-existing, antigen-specific memory B-cell clones, thereby enabling the generation of antigen-specific hybridomas even in the absence of FDCs. Alternatively, successful engraftment of FDCs into this humanized mouse model may enhance affinity maturation and promote a more efficient production of antigen-specific antibodies, an avenue of inquiry that warrants further investigation.

The limited human B cell repertoire remains a major challenge for improvement in our humanized mouse model. Because the number of B-cell clones gradually declined over the 3-month study period, peptide immunization was initiated on the day of PBMC transplantation. Given that approximately 5×10L PBMCs were administered per mouse, the total number of functional B cells, especially those with epitope-specific repertoires, may be insufficient. Epitope-specific B cells are estimated to occur at a frequency of 1 in 300,000, whereas protein-specific B cells are estimated at approximately 1 in 40,000 (43). Since B cells constitute about 10% of PBMCs, fewer than 500,000 B cells were likely transferred per mouse, implying that each mouse may have received only a single epitope-specific B cell clone. Nevertheless, most mice in our study successfully produced antigen-reacting antibodies against CH401 or BAL6MAP epitopes, and the number of peptide-reactive clones increased during hybridoma screening. If a full-length protein antigen containing multiple epitopes had been used instead of a single 20-mer peptide, a greater number of antigen-reacting B cell clones might have been generated. In contrast, T cells exhibit approximately tenfold greater repertoire diversity than B cells, allowing their responses to arbitrary epitopes easier to detect and analyze. The limited diversity of the B cell repertoire, however, constrains its ability to respond to novel antigens in this system.

Several clones increased in the mice, appearing to have expanded without immunization. Since these clones were not observed in the original PBMCs, they were likely expanded independently in each mouse. The antigens may have originated from the mice themselves or from environmental microbes. The expansion of a single clone suggests that stable clonal expansion may be possible in this mouse system, which should be addressed in future studies.

A key observation from our previous studies was the markedly reduced incidence of GVHD in NOG-hIL-4-Tg mice compared with NOG mice under most transplantation conditions. This led us to hypothesize that elevated plasma hIL-4 levels may suppress the onset of this life-threatening immune reaction (22). In the present study, optimal plasma hIL-4 concentrations in NOG-hIL-4-Tg mice ranged between 100 pg/mL and 500 pg/mL and showed a positive linear correlation with the number of antigen-specific B cell clones and Ig class-switching events. IL-4 has been reported to protect naïve B cells from apoptosis by upregulating BCL-xL (44) and to exert different effects on B cells at various developmental stages (45). Naïve B cells may increase the expression of the transcription factor BCL-6, which in turn enhances their survival. However, IL-4 negatively regulates BCL-6 expression in GC B cells, thereby suppressing the accumulation of fully differentiated plasma cells. Although B cell expansion in the mouse system may be attributable to one or more of these IL-4–related mechanisms, the reason why human plasmablasts accumulated in the spleen and bone marrow at the expense of long-lived plasma cells remains unresolved.

In addition, the likelihood that elevated hIL-4 had enhanced the induction of activation-induced cytidine deaminase (AID) is consistent with previous reports (46). Furthermore, hIL-4 may suppress glucocorticoid receptor expression in B cells, potentially reducing their susceptibility to stress-induced apoptosis (25). Although we were unable to assess AID expression in B cells from engrafted mice due to insufficient cell numbers, this potential regulatory mechanism warrants further investigation. An important limitation of this system is that long-term survival of human B cells or plasma cells was not maintained beyond 3 months, during which human T cells became dominant. The factors required to sustain B cell longevity will be investigated in future studies.

Despite the presence of optimal hIL-4 levels, some humanized mice (2 of 24; Supplementary Table 1) still developed GVHD during the 3 months of observation. This finding suggests that although elevated hIL-4 may delay or mitigate GVHD severity in most cases, it does not fully prevent its onset. The observed reduction in human B and T cell clone numbers to approximately 50% may reflect the expansion of mouse-specific clones within the humanized mouse environment and/or the loss of human clones due to insufficient survival signals, although most mice remained viable and generated antigen-specific responses enabling additional in vivo analysis of human adaptive immunity. The effects of IL-4 on B cell survival and differentiation may contribute to the delay of GVHD, as B cells can suppress T cell function through the production of IL-10, an immunosuppressive cytokine (47). B cell activation via STAT-3 is known to induce IL-10 secretion (48). Because B cells have a short lifespan, the loss of engrafted B cells may enhance T cell activation and inflammation, leading to the subsequent development of GVHD. Therefore, improving the lifespan of human B cells in this system may help to reduce or prevent the occurrence of GVHD.

Based on our findings and previous reports, hIL-4 expression is effective for evaluating antibody responses in PBMC-transplanted humanized mice. However, the NOG-hIL-4-Tg model has some inherent limitations that affect its utility for fundamental immunological studies. These include incomplete maintenance of naïve lymphocytes, impaired memory cell formation, particularly within the B cell lineage, absence of these cells in the lymph node or peritoneal cavity, and an overall restricted immune reconstitution such as the development of macrophages and NK cells, and lack of germinal center formation and lymph nodes. Furthermore, the potential onset of GVHD complicates data interpretation and may limit the feasibility of experiments that require long-term observation. Although the limited diversity and persistence of the human B cell repertoire remain significant challenges, these limitations may be partially overcome by increasing the number of transplanted B cells. Our future efforts will now focus on promoting memory B cell formation and elucidating the mechanisms underlying germinal center development to enhance the model’s capacity for affinity maturation of B cell clones and to more accurately recapitulate human adaptive immunity.

Recently, Chupp et al. (49) reported a new humanized THX mouse model in which oral estradiol treatment for at least four weeks enabled successful engraftment of human cord blood CD34L cells. This model reconstituted a functional human lymphoid and myeloid immune system with diverse B cell and T cell antigen receptor repertoires, which mounted mature T cell–dependent and –independent antibody responses, including somatic hypermutation, class-switch recombination, and plasma cell and memory B cell differentiation. Therefore, this model may be suitable for studies requiring long-term or fully mature antibody responses. However, in HSC-transplantation systems, including those not limited to the Chupp group, immune cells develop within the murine environment. Consequently, the established immune system and its composition do not fully reflect those of the donor. Moreover, because the immune environment of cancer patients differs substantially from that of healthy donors, humanized mice reconstructed with HSCs cannot faithfully reproduce the immune context of cancer patients for drug evaluation. To establish donor-specific immune environments and compare humoral immune responses between donors—such as healthy individuals and cancer patients—PBMCs from patients may be more appropriate. Although our PBMC-based humanized mouse system has not yet been fully optimized for germinal center formation, lymph node development, or robust affinity maturation, it does support the survival of human T and B cells, maintain donor-derived lymphocytes, and activate them in response to immune drugs with the potential to engage the donor’s immune system. Estradiol treatment may also be beneficial in our system to further optimize the reconstitution of a proper human lymphoid and myeloid immune system.

In conclusion, limited concentrations of IL-4 enhance human B cell survival and IgG production in this xenograft model. Our study suggests the potential utility of this humanized mouse system for preliminary or short-term in vivo evaluation of vaccine responses, antibody generation, and immunotherapies.

## Author Contributions

Yoshie Kametani and RI contributed to the conception and design of this study. RI and Yoshie Kametani contributed to methodology development. SO, YO, SY, YH, AM, MS, NK, MM, HK, DK, Tomoka Shimizu, MK, Yusuke Kikuchi, SN, RO and YN contributed to the data acquisition and analysis. Yoshie Kametani, JKK, TM, HI and Takashi Shiina contributed to the writing, reviewing, and revision of the manuscript., RI, AY, AH, TS, and BT provided administrative, technical, and material support. Takashi Shiina and Yoshie Kametani supervised the study. All the authors contributed to the manuscript and approved the submitted version.

## Acknowledgments

We thank the members of the Teaching and Research Support Center at Tokai University School of Medicine for their technical skills.

