## Supplemental Tables for "Human Interleukin-4-Dependent Facilitation of Human IgG Production in PBL-NOG-hIL-4-Tg mice": Supplemental Tables 3-2.pdf

### Supplemental Table 1 List of donors and mice used in the repertoire analysis

Donor, donor ID; Weeks, period of engraftment; mouse ID, mouse ID for repertoire analysis; IL-4, hIL-4 concentration in mouse plasma (pg/ml); sex, mouse sex; SPL cell, number of mouse spleen cells. Pink columns represent mice with low plasma IL-4 levels. Gray columns represent mice that died before completion of the analyses.

| Donor | Weeks | mouse # | IL-4 | sex | SPL cell |
|---|---|---|---|---|---|
| RG1 | ① 4W | TIK102 | 235.651 | ♀ | 1.1E+08 |
| | | TIK103 | 274.057 | ♂ | 9.9E+06 |
| | ② 8W | TIK104 | 135.180 | ♀ | 7.1E+07 |
| | | TIK105 | 724.625 | ♂ | 2.5E+07 |
| | ③ 12W | TIK106 | 213.018 | ♀ | 1.1E+07 |
| | | TIK107 | 170.049 | ♂ | 2.1E+05 |
| RG2 | ① 4W | TIK109 | 270.049 | ♀ | 9.7E+06 |
| | | TIK110 | 15.551 | ♀ | 4.9E+07 |
| | ② 8W | TIK1111 | 255.725 | ♀ | 8.0E+07 |
| | | TIK112 | 21.700 | ♂ | 2.5E+05 |
| | ③ 12W | TIK113 | 186.837 | ♂ | 3.1E+06 |
| | | TIK114 | 17.657 | ♀ | |
| RG3 | ① 4W | TIK116 | 217.102 | ♀ | 5.7E+06 |
| | | TIK117 | 260.790 | ♂ | 7.3E+06 |
| | ② 8W | TIK118 | 232.164 | ♂ | 7.3E+07 |
| | | TIK119 | 302.358 | ♂ | 8.7E+07 |
| | ③ 12W | TIK120 | 267.421 | ♂ | 3.5E+07 |
| | | TIK121 | 234.594 | ♀ | |
| RG4 | ① 4W | TIK123 | 252.404 | ♂ | 1.1E+07 |
| | | TIK124 | 276.554 | ♀ | 3.3E+07 |
| | ② 8W | TIK125 | 174.965 | ♂ | 8.4E+07 |
| | | TIK126 | 207.664 | ♂ | 6.5E+07 |
| | ③ 12W | TIK127 | 182.450 | ♂ | 6.7E+07 |
| | | TIK128 | 207.503 | ♂ | |
| | (gray) | Dead before analysis | | | |
| | (pink) | IL-4 low concentration | | | |

#### Supplemental Table 2 Antibodies used for FCM and IHC

| Antibody | Clone | Company | Cat.No. |
|---|---|---|---|
| for FCM | | | |
| FITC anti-human CD3 | UCHT1 | BioLegend | 300406 |
| APC anti-human CD4 | RPA-T4 | BioLegend | 300514 |
| PE/Cy7 anti-human CD5 | UCHT2 | BioLegend | 300622 |
| Alexa Fluor®700 anti-human CD8 | HIT8a | BioLegend | 300920 |
| PerCP/Cy5.5 anti-human CD11c | Bu15 | BioLegend | 337209 |
| APC/Cy7 anti-human CD19 | HIB19 | BioLegend | 302218 |
| PE anti-human CD25 | BC96 | BioLegend | 302606 |
| PE anti-human CD27 | M-T271 | BD Bioscience | 555441 |
| Alexa Fluor®700 anti-human CD38 | HIT2 | BioLegend | 303524 |
| Pacific Blue™ anti-human CD45 | HI30 | BioLegend | 304022 |
| PE anti-human CD163 | GHI/61 | BioLegend | 333605 |
| PerCP/Cy5.5 anti-human CD279 (PD-1) | EH12.2H7 | BioLegend | 329914 |
| FITC anti-human HLA-DR | LN3 | eBioscience | 11-9956-73 |
| PI | | SIGMA | P4170 |
| For IHC | | | |
| CD3 | PS1 | NICHIREI BioSciences | 413241 |
| CD4 | 1F6 | NICHIREI BioSciences | 413181 |
| CD8 | 4B11 | Leica Biosystems | NCL-L-CD8-4B11 |
| CD20 | L26 | NICHIREI BioSciences | 422441 |

### Supplemental Table 3 Statistical analysis of specific clone numbers

**Donor; the donor ID, Weeks; the period of engraftment, mouse ID; the mouse ID used for the repertoire analysis, TCRA; TCR$\alpha$ chain, TCRB; TCR$\beta$ chain, IgM; IgM chain, IgG; IgG chain, IgA; IgA chain, IgK; Ig$\kappa$ chain, IfL; Ig$\lambda$ chain. PBMC; original PBMCs used for the transplantation. 4W, 8W, 12W; spleen cells obtained after 4 weeks, 8 weeks and 12 weeks. Average and SD were shown in each group. One way-ANOVA and Student's *t*-test were conducted and the *t*-values are shown. Yellow markers show the *t*-values larger than p=0.05. Pink marker indicates the Low hIL-4 mouse.**

| | mouse ID | Number of clones | | | | | | |
|---|---|---|---|---|---|---|---|---|
| | | TRA | TRB | IgM | IgG | IgA | IgK | IgL |
| PBMC | TIK101 | 24778 | 12626 | 39618 | 15644 | 32920 | 13293 | 22989 |
| | TIK108 | 33221 | 11971 | 39215 | 16276 | 26379 | 15182 | 20509 |
| | TIK115 | 21448 | 10328 | 27362 | 11812 | 13157 | 11872 | 16375 |
| | TIK122 | 24344 | 20590 | 35619 | 21718 | 28673 | 18891 | 26607 |
| AVERAGE | | 25947.75 | 13878.75 | 35453.5 | 16362.5 | 25282.25 | 14809.5 | 21620 |
| SD | | 4389.995579 | 3964.13909 | 4924.21834 | 3532.42675 | 7383.3759 | 2632.77786 | 3724.53608 |
| 4W | TIK102 | 10250 | 4569 | 31036 | 17654 | 24118 | 7653 | 12588 |
| | TIK103 | 14688 | 6282 | 26517 | 20211 | 16969 | 7310 | 9031 |
| | TIK109 | 9764 | 3249 | 25483 | 25206 | 20473 | 10404 | 12590 |
| | TIK110 | 10091 | 9359 | 24598 | 19672 | 11916 | 6602 | 7304 |
| | TIK116 | 11119 | 10355 | 18063 | 14713 | 12858 | 5054 | 6474 |
| | TIK117 | 14022 | 6397 | 22120 | 17222 | 14003 | 6851 | 6871 |
| | TIK123 | 10906 | 7010 | 26896 | 19128 | 18998 | 10128 | 10958 |
| | TIK124 | 12890 | 7957 | 29603 | 24835 | 22586 | 9822 | 12951 |
| AVERAGE | | 11716.25 | 6897.25 | 25539.5 | 19830.125 | 17740.125 | 7978 | 9845.875 |
| SD | | 1772.755253 | 2194.06271 | 3840.56946 | 3397.32724 | 4265.07566 | 1808.77866 | 2580.51998 |
| 8W | TIK104 | 8571 | 4173 | 15091 | 10167 | 18631 | 5957 | 7795 |
| | TIK105 | 7248 | 6083 | 18949 | 10974 | 12392 | 4726 | 4688 |
| | TIK111 | 9186 | 5132 | 31540 | 12701 | 14419 | 9570 | 11095 |
| | TIK112 | 11951 | 8414 | 10084 | 11933 | 6731 | 6698 | 7421 |
| | TIK118 | 12756 | 9041 | 17970 | 12345 | 10625 | 3440 | 7039 |
| | TIK119 | 10106 | 5799 | 32213 | 14404 | 16080 | 6477 | 8297 |
| | TIK125 | 6853 | 5516 | 34725 | 9292 | 9178 | 4427 | 9859 |
| | TIK126 | 10707 | 4847 | 22524 | 22087 | 14120 | 6107 | 6932 |
| AVERAGE | | 9672.25 | 6125.625 | 22887 | 12987.875 | 12772 | 5925.25 | 7890.75 |
| SD | | 1975.962155 | 1606.67093 | 8415.00042 | 3742.99935 | 3604.94424 | 1734.90762 | 1817.91039 |
| 12W | TIK106 | 6201 | 4477 | 18025 | 11328 | 15040 | 7525 | 7576 |
| | TIK107 | 7723 | 4498 | 5577 | 4953 | 4862 | 1739 | 714 |
| | TIK113 | 9833 | 7049 | 13319 | 7336 | 5535 | 4925 | 4658 |
| | TIK120 | 8901 | 6494 | 19000 | 12714 | 11416 | 4715 | 5600 |
| | TIK127 | 16898 | 9655 | 38382 | 17665 | 24058 | 9886 | 8267 |
| AVERAGE | | 9911.2 | 6434.6 | 18860.6 | 10799.2 | 12182.2 | 5758 | 5363 |
| SD | | 3698.534759 | 1914.92084 | 10853.2583 | 4410.37656 | 7036.46651 | 2760.31781 | 2664.46918 |
| ANOVA(1way) | | 1.37757E-07 | 0.00042273 | 0.03708318 | 0.00340995 | 0.00812321 | 2.1243E-05 | 1.5762E-07 |
| Turkey HSD | PBMC vs 4W | *P<0.05 | *P<0.05 | P>0.05 | P>0.05 | P>0.05 | *P<0.05 | *P<0.05 |
| | PBMC vs 8W | *P<0.05 | *P<0.05 | P>0.05 | P>0.05 | *P<0.05 | *P<0.05 | *P<0.05 |
| | PBMC vs 12W | *P<0.05 | *P<0.05 | *P<0.05 | P>0.05 | *P<0.05 | *P<0.05 | *P<0.05 |
| | 4W vs 8W | P>0.05 | P>0.05 | P>0.05 | *P<0.05 | P>0.05 | P>0.05 | P>0.05 |
| | 4W vs 12W | P>0.05 | P>0.05 | P>0.05 | *P<0.05 | P>0.05 | P>0.05 | P>0.05 |
| | 8W vs 12W | P>0.05 | P>0.05 | P>0.05 | P>0.05 | P>0.05 | P>0.05 | P>0.05 |

### Supplemental Table 4 Representative data of class-switched clones in a NOG-hIL-4-Tg mouse

Representative data of class-switched clones in a NOG-hIL-4-Tg mouse
V gene, variable gene; D gene, diversity gene; J gene, joining gene; CDR sequence, complementarity determining region sequence; Sample, TIK101hA; original PBMC of HD1 IgA, TIK101hG; IgG, TIK101hM; IgM, TIK102hA; IgA, TIK102hG; IgG, TIK102hM; IgM. The percentage of the clones per total clones of each type is shown.

Please find in the attached Excel file.

#### Supplemental Table 5. Antigen-reactive polyclonal antibodies in the NOG-hIL-4-Tg mice

PBS, CH401MAP or BAL6MAP was immunized into NOG-hIL-4-Tg mice. mice (n=6 for PBS, n=6 for CH401MAP and n=4 for BAL6MAPThe IL-4 concentration in the sera and titer of the plasma were shown.

| Immunogen | Mouse # | IL-4 (ng/ml) | anti-CH401 (ng/ml) | anti-BAL6 (ng/ml) |
|---|---|---|---|---|
| PBS | #563 | 312.0 | 28.6 | 3.9 |
| | #565 | 312.0 | 6.2 | 1.1 |
| | #923 | 235.3 | 62.7 | 61.0 |
| | #925 | 188.2 | 3.2 | 6.4 |
| | #937 | 182.0 | 43.4 | 38.4 |
| | #948 | 281.7 | 2.6 | 1.9 |
| CH401MAP | #487 | 300.6 | 10.4 | 1.4 |
| | #511 | 267.6 | 8.1 | 2.5 |
| | #539 | 201.1 | 19.7 | 5.2 |
| | #271 | 283.3 | 11.7 | nd |
| | #405 | 163.6 | 0.9 | nd |
| | #419 | 194.0 | 15.8 | nd |
| BAL6MAP | #915 | 296.1 | 21.3 | 22.6 |
| | #916 | 273.4 | 12.2 | 12.7 |
| | #928 | 148.4 | 93.2 | 42.3 |
| | #929 | 192.0 | 69.9 | 32.8 |
