## Supplemental Figures for "Human Interleukin-4-Dependent Facilitation of Human IgG Production in PBL-NOG-hIL-4-Tg mice"

### Human Interleukin-4-dependent facilitation of anti-peptide Ig production with antibody class switching and diverse B and T cell repertoires

Yoshie Kametani[1,2*], Shino Ohshima[1], Ryoji Ito[3], Yusuke Ohno[3], Soga Yamada[1], Yuki Hoshino[1], Asuka Miyamoto[1], Mao Suzuki[1], Nagi Katano[1] Banri Tsuda[4], Mariko Miyazawa[5] , Hirofumi Kashiwagi[5], Daiki Kirigaya[1], Tomoka Shimizu[1], Mika Kojima[1], Yusuke Kikuchi[1], Shunsuke Nakada[1], Rentaro Ohki[1], Atsushi Yasuda[6], Ayako Hirota[7], Toshiro Seki[6], Yukio Nakamura[8] , Jerzy K. Kulski[1,9] , Tomotaka Mabuchi[7], Hitoshi Ishimoto[5], and Takashi Shiina[1,2]

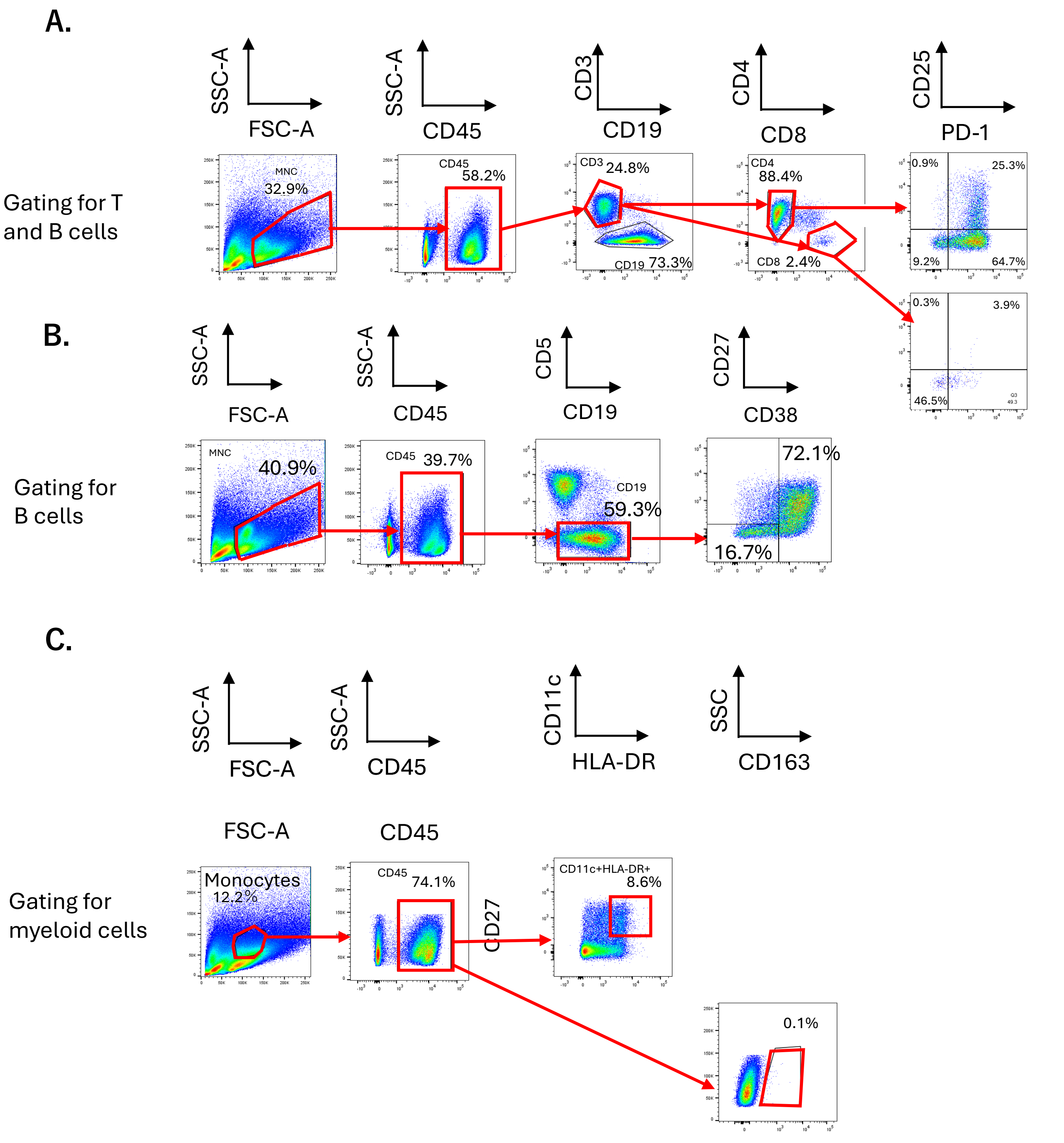


#### Supplemental Figure 1 Lymphocyte gating

(A) Gating strategy for T cell subsets and B cells with activation markers in PBL-NOG-hIL-4-Tg spleen cells. Cells were gated within the mononuclear cell gate (FSC: 80K–250K). (B) Gating strategy for distinguishing plasmablasts among B cells in PBL-NOG-hIL-4-Tg spleen cells. Cells were gated within the mononuclear cell gate (FSC: 80K–250K). Red squares: Gated cell populations. Red arrows: Indicate the next gating steps. (C) Gating strategy for dendritic cells (DCs) and macrophages in PBL-NOG-hIL-4-Tg spleen cells. Red squares: Gated cell populations. Red arrows: Indicate the next gating steps.

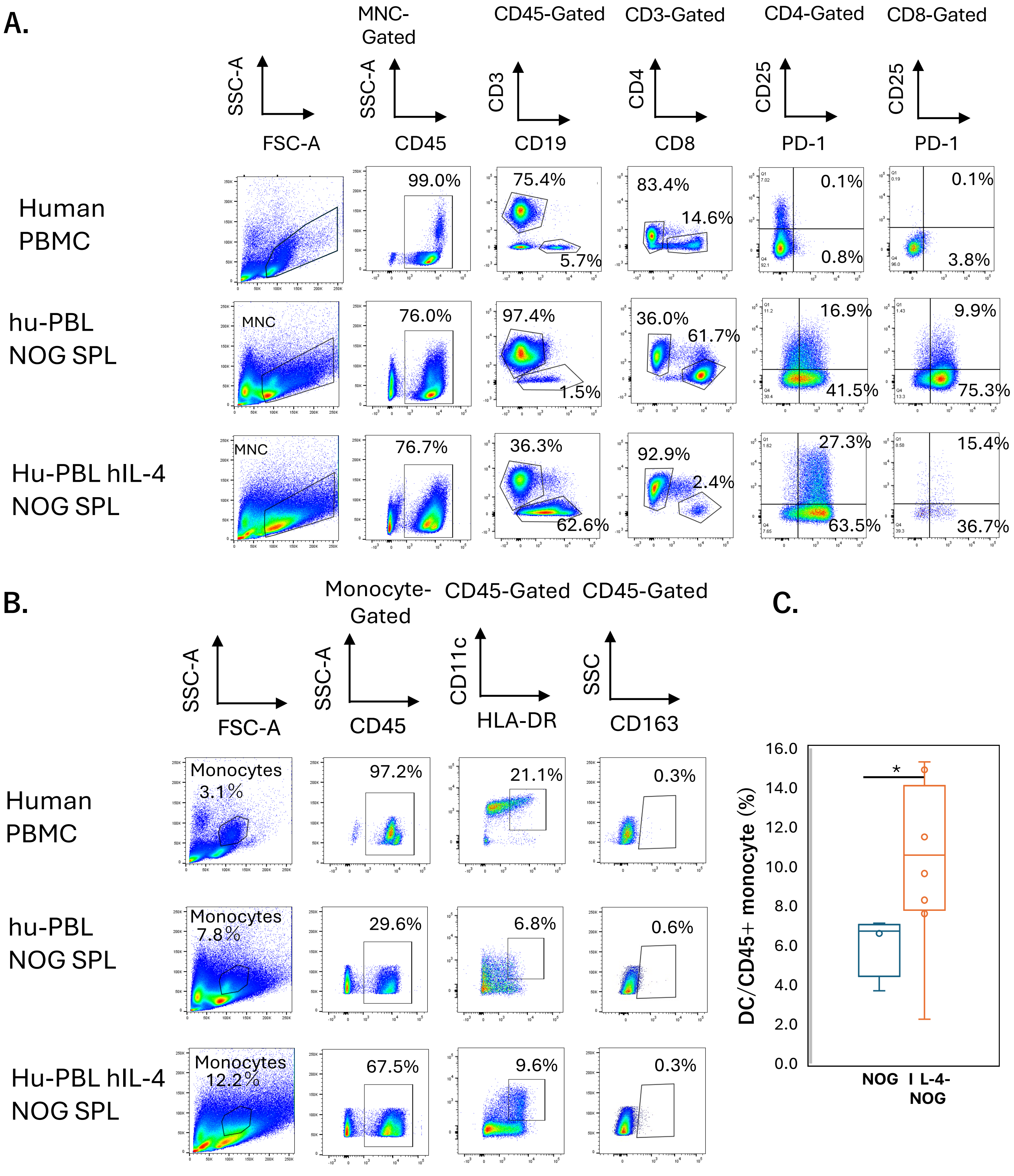


**Supplemental Figure 2 Flow cytometry of engrafted T Cells and myeloid cells in hu-PBL hIL-4 NOG mice.**

**(A) Representative flow cytometry (FCM) patterns of lymphocyte-gated PBMCs and mouse spleen cells. Cells were further gated for CD45⁺ populations, T cell markers, and activation/exhaustion markers.**

**(B) Monocyte-gated PBMCs and mouse spleen cells were analyzed for CD45⁺ populations. DCs were identified as CD11c⁺HLA-DR⁺ cells, and macrophages as CD163⁺ cells. Upper panels: Original PBMCs. Middle panels: Spleen cells from NOG-engrafted mice. Lower panels: Spleen cells from NOG-hIL-4-Tg-engrafted mice. The percentages of gated subsets are shown in each panel. (C) Mean percentage of DCs in NOG versus NOG-hIL-4-Tg spleens (IL-4-NOG). NOG (n = 4), NOG-hIL-4-Tg (n = 8).Student's *t*-test was performed, with significance indicated as follows: $P < 0.05$ (*).**

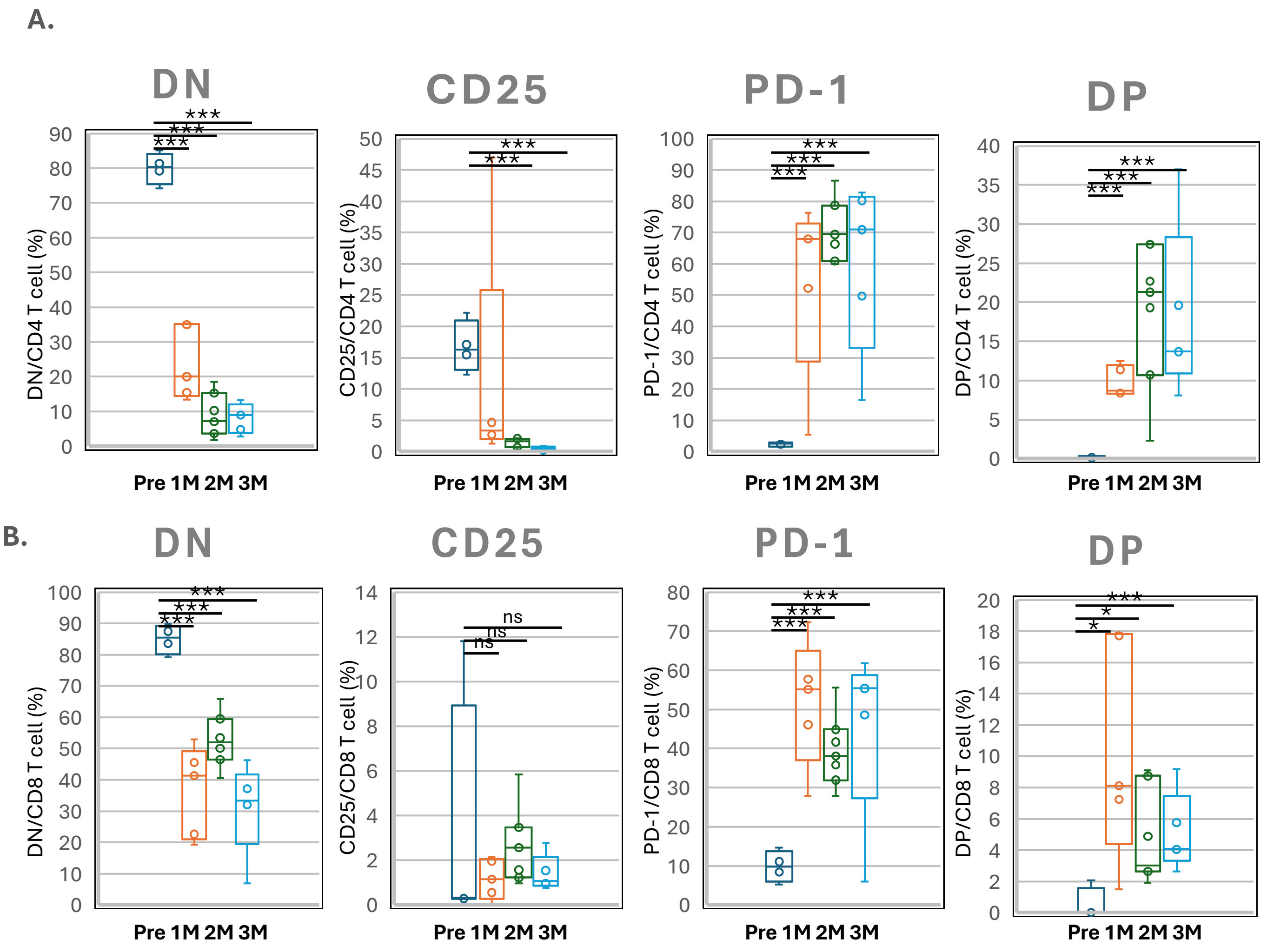


**Supplemental Figure 3. CD25 and PD-1 expression on engrafted human T cells in NOG-hIL-4-Tg mice.**

**(A) $CD4^+$ T cells, (B) $CD8^+$ T cells. Double-negative (DN): $CD25^-PD\text{-}1^-$ double-negative T cells, CD25: $CD25^+$ single-positive T cells, PD-1: $PD\text{-}1^+$ single-positive T cells, Double-positive (DP): $CD25^+PD\text{-}1^+$ double-positive T cells. Time points: Pre ($n = 4$), 1M ($n = 7$), 2M ($n = 7$), 3M ($n = 3$). TIK107 and TIK113 were not analyzed due to insufficient cell numbers for flow cytometry. Data are presented as mean ± S.D. Student's *t*-test was performed, with significance indicated as follows: $P < 0.05$ (*), $P < 0.001$ (**), ns = not significant. Sample details are provided in Table 1B.**

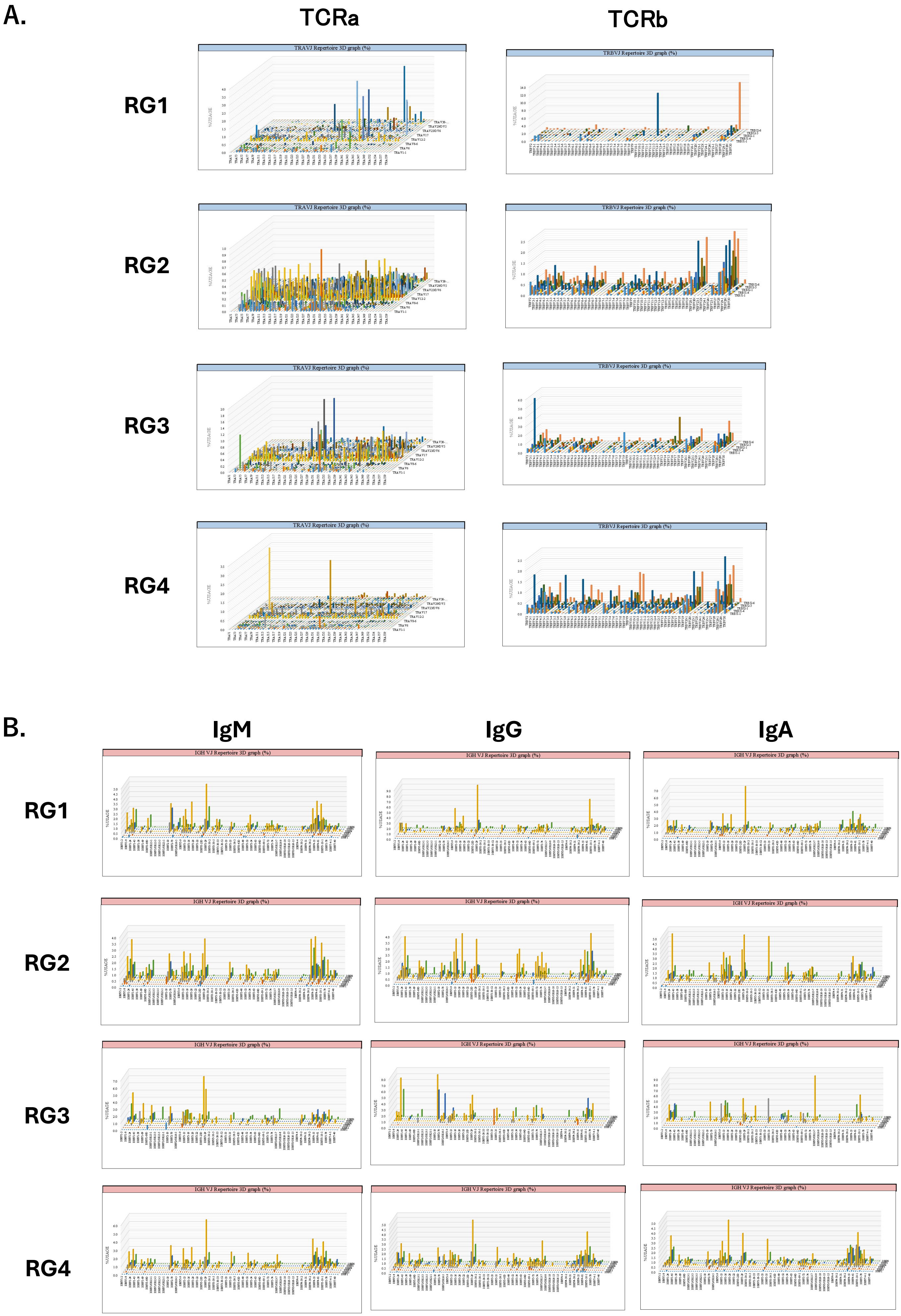
A.
TCRa
TCRb
RG1
RG2
RG3
RG4
TRAVJ Repertoire 3D graph (%)
TRBVJ Repertoire 3D graph (%)
%USAGE
B.
IgM
IgG
IgA
IGH VJ Repertoire 3D graph (%)

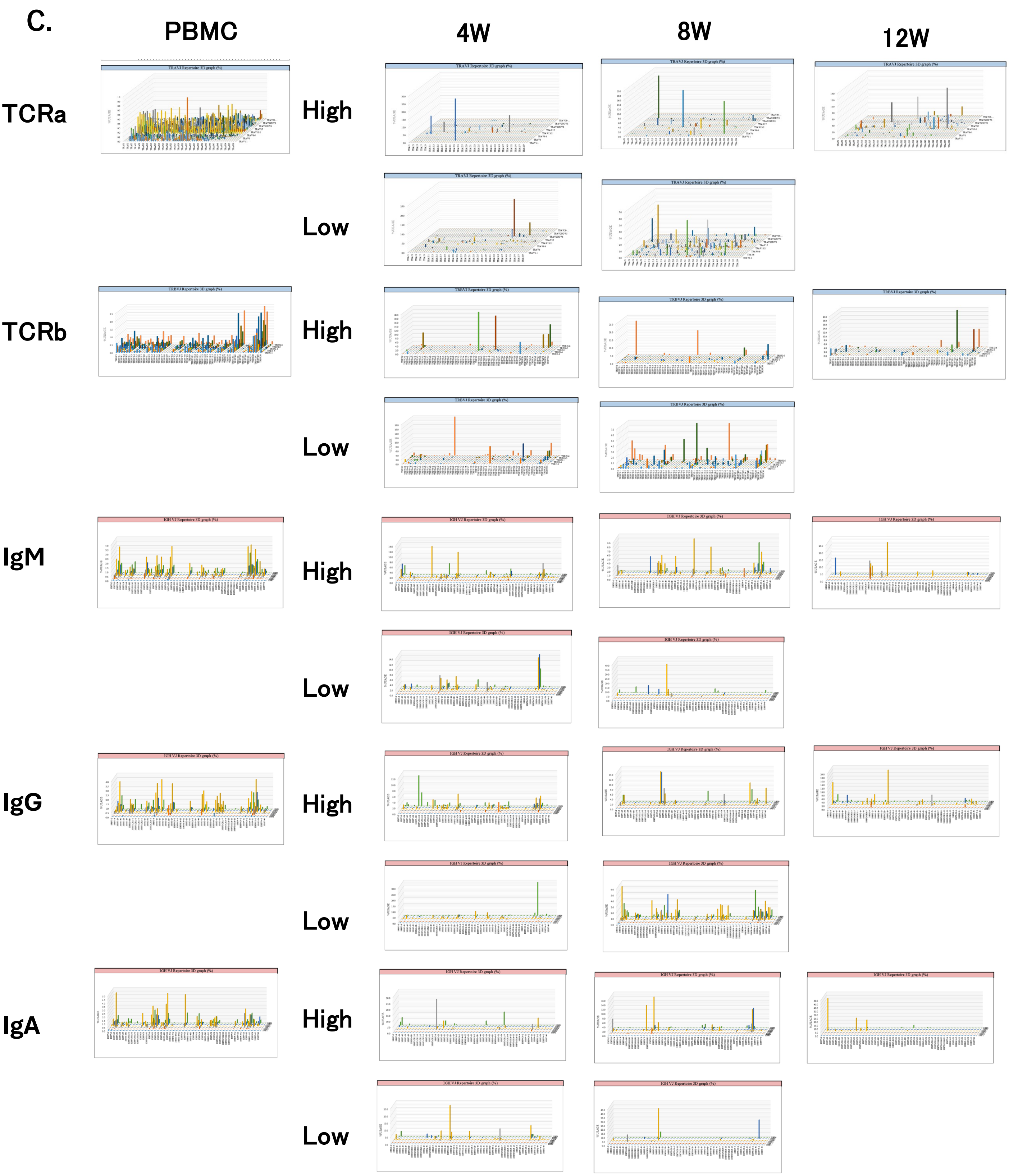


Supplemental Figure 4. TCR and Ig repertoire of individual donor and mouse

(A) TCRa(left panels) and TCRb(right panels) repertoire of RG1, RG2, RG3 and RG4 PBMCs. Vertical lines are numbers of clones, and horizontal lines of two directions are repertoires. (B) IgM (left panels), IgG (middle panels) and IgA (right panels) repertoire of RG1, RG2, RG3 and RG4 PBMCs. (C) Representative RG2 repertoire profile of original PBMC, mouse spleen cells of 4 weeks, 8 weeks and 12 weeks after the transplantation. Upper panels (High) are the mice with normal IL-4 concentration. Lower panels (Low) are the mice with lower IL-4 concentration. Detail was shown in Supplementary Table 1

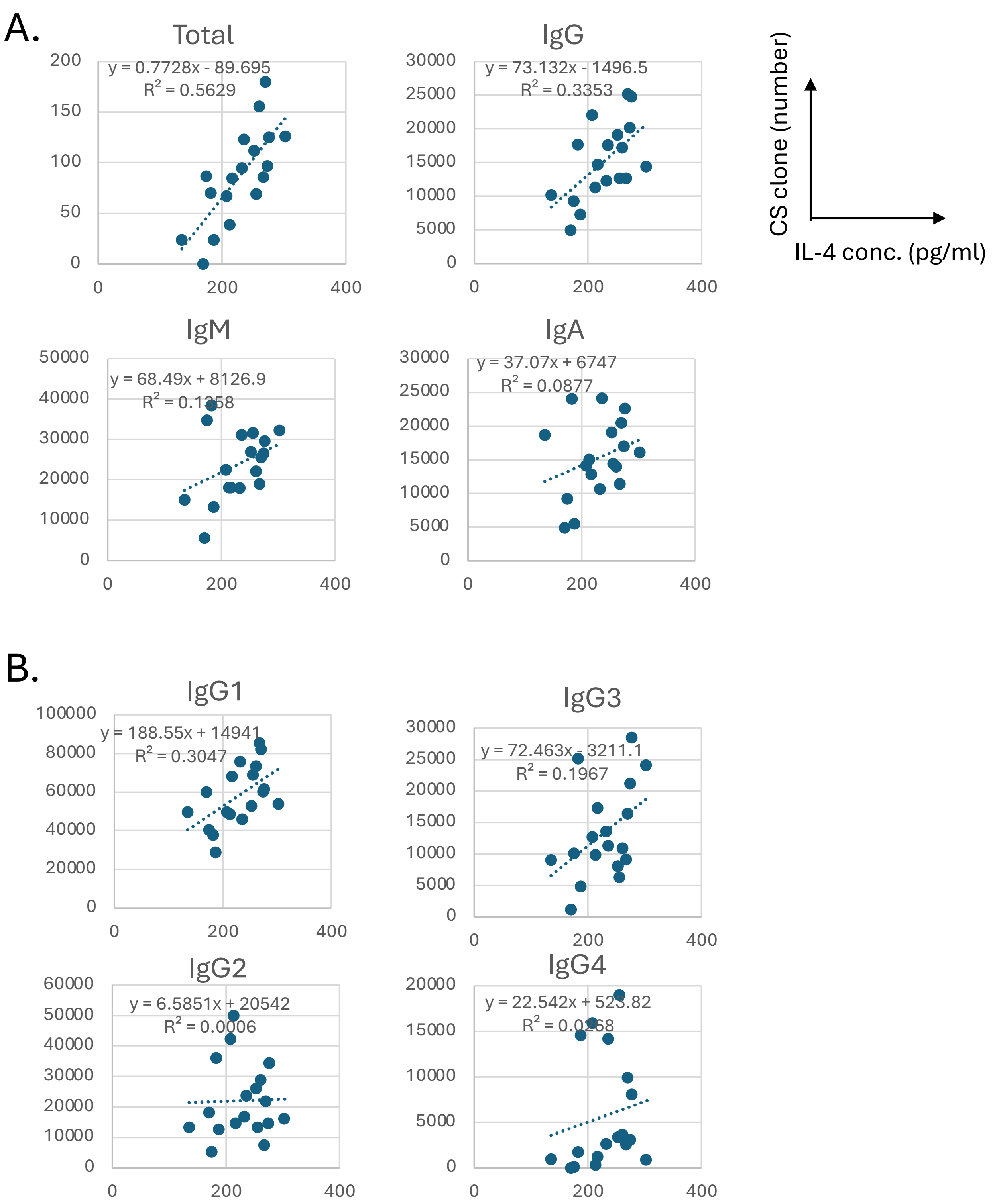


#### Supplemental Figure 5. Correlation between IL-4 concentration and the number of class-switched clones

(A) Total clones (>0.01% of total), IgM, IgG, IgA.
(n = 18). Samples with low IL-4 levels (<100 pg/mL; TIK110 and TIK112) and high IL-4 levels (>500 pg/mL; TIK105) were excluded. The regression lines, formulas, and $R^2$ values are displayed in each panel. F-values for each marker: Total clones = 0.0003349, IgM = 0.1323, IgG = 0.0118, IgA = 0.2329.
(B) Total clones (>0.01% of total), (B) IgM, (C) IgG, (D) IgA.
(n = 18). Samples with low IL-4 levels (<100 pg/mL; TIK110 and TIK112) and high IL-4 levels (>500 pg/mL; TIK105) were excluded. The regression lines, formulas, and $R^2$ values are displayed in each panel. F-values for each marker: IgG1= 0.017, IgG2 =0.9229, IgG3 = 0.0653Ig4 = 0.5159.

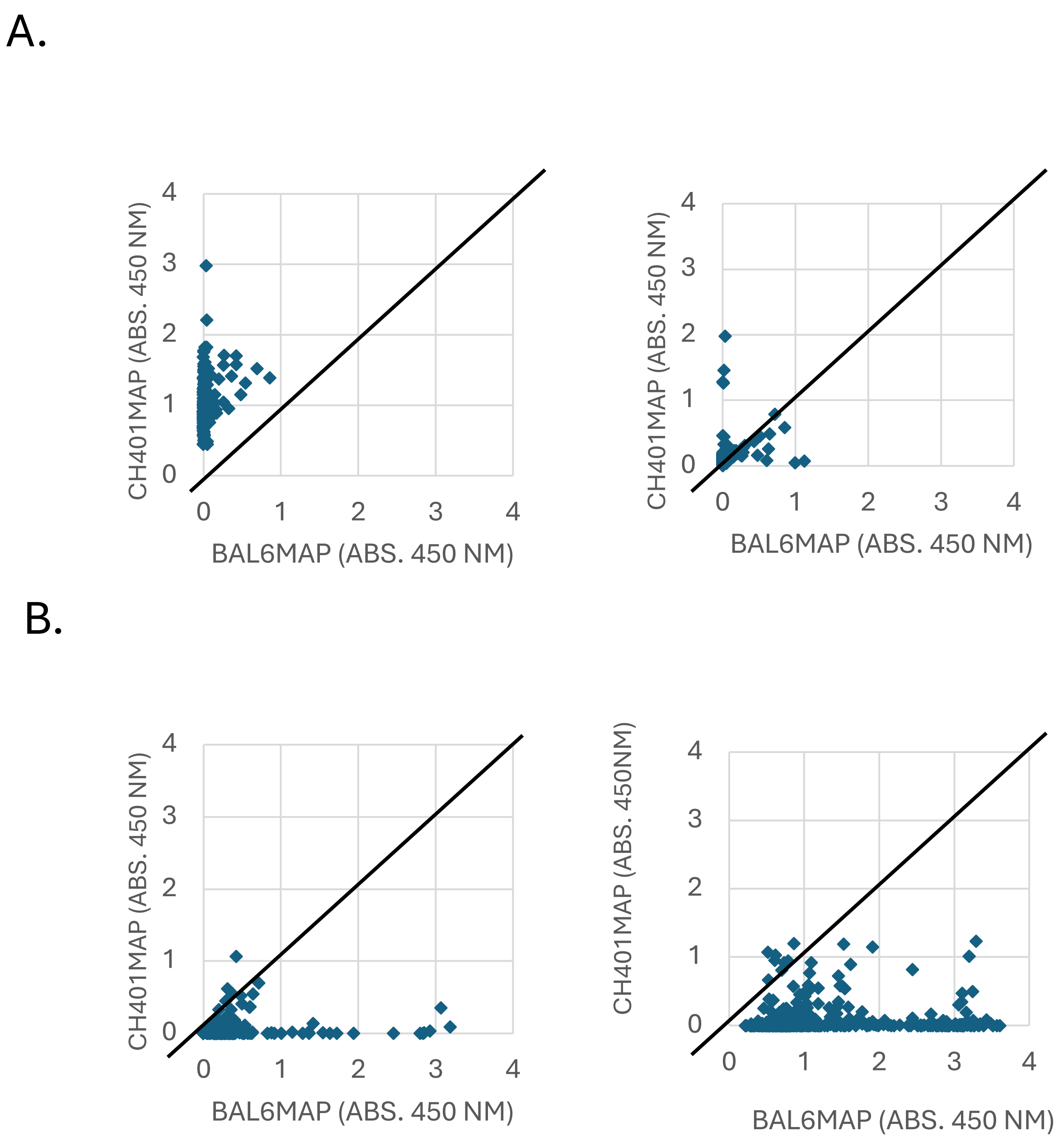


**Supplemental Figure 6. Antigen-specific clones produced by BALB/c mice**

(A) CH401MAP-immunized mouse clones (n = 2). Left panel: BALB1, Right panel: BALB2. The antibody titer for both mice was greater than 1:1,000.
(B) BAL6MAP-immunized mouse clones (n = 2). Left panel: BALB3, Right panel: BALB4. The antibody titer for both mice was greater than 1:100,000. Each dot represents a single clone.
